# scJoint: transfer learning for data integration of atlas-scale single-cell RNA-seq and ATAC-seq

**DOI:** 10.1101/2020.12.31.424916

**Authors:** Yingxin Lin, Tung-Yu Wu, Sheng Wan, Jean Y.H. Yang, Wing H. Wong, Y. X. Rachel Wang

**Author notes:** Equal contribution (Yingxin Lin, Tung-Yu Wu). To whom correspondence should be addressed. Email: Y. X. Rachel Wang,; Wing H. Wong,.

## Abstract

Single-cell multi-omics data continues to grow at an unprecedented pace, and effectively integrating different modalities holds the promise for better characterization of cell identities. Although a number of methods have demonstrated promising results in integrating multiple modalities from the same tissue, the complexity and scale of data compositions typically present in cell atlases still pose a significant challenge for existing methods. Here we present scJoint, a transfer learning method to integrate atlas-scale, heterogeneous collections of scRNA-seq and scATAC-seq data. scJoint leverages information from annotated scRNA-seq data in a semi-supervised framework and uses a neural network to simultaneously train labeled and unlabeled data, enabling label transfer and joint visualization in an integrative framework. Using multiple atlas data and a biologically varying multi-modal data, we demonstrate scJoint is computationally efficient and consistently achieves significantly higher cell type label accuracy than existing methods while providing meaningful joint visualizations. This suggests scJoint is effective in overcoming the heterogeneity in different modalities towards a more comprehensive understanding of cellular phenotypes.

## Introduction

Advances in single-cell technologies have enabled comprehensive studies of cell heterogeneity, developmental dynamics, and cell communications across diverse biological systems at an unprecedented resolution. There are a variety of protocols profiling the transcriptomics, as exemplified by single-cell RNA-seq (scRNA-seq). In addition, a number of technologies have been developed for other molecular measurements in individual cells towards building a more holistic view of cell functions, including chromatin accessibility, protein abundance, and methylation [1].

In particular, single-cell ATAC-seq (scATAC-seq) is an epigenomic profiling technique for measuring chromatin accessibility to discover cell type specific regulatory mechanisms [2, 3]. scATAC-seq offers a complementary layer of information to scRNA-seq, and together they provide a more comprehensive molecular profile of individual cells and their identities. However, it has been noted that the extreme sparsity of scATAC-seq data often limits its power in cell type identification [4]. In contrast, large amounts of well-annotated scRNA-seq datasets have been curated as cell atlases [5, 6], motivating us to transfer cell type information from scRNA-seq to scATAC-seq for better classification of cell types in an integrative analysis framework.

A number of methods exist to denoise, batch correct, and perform integration of single-omics data across multiple experiments for both transcriptomic data [7–12] and scATAC-seq data [13]. However, direct applications of these methods to multi-omics data integration are computationally challenging and often suboptimal, since different modalities have vastly different dimensions and sparsity levels. Recently, a growing number of methods have been proposed to address the need for integrative analysis across different modalities. When the data consist of simultaneous multi-modal measurements within the same cell [14, 15], methods like scAI [16] and MOFA+ [17] have been developed based on factor analysis and joint clustering. In general, these paired measurements are technically more challenging and costly to perform.

More commonly, different modalities are derived from different cells taken from the same or similar populations. In this setting, most existing methods can be broadly divided into four categories: manifold alignment [18–20], matrix factorization (Liger [21], coupledNMF [22]), using correlations to identify nearby cells across modalities (Conos [23], Seurat [24]), or neural network approaches, each with its own limitations when facing complex data compositions as typically seen in cell atlases. Manifold alignment methods have demonstrated promising results in integrating multiple modalities from the same tissue. However, requiring distributions to match globally is often too restrictive when different modalities are derived from different tissues and cell types. Furthermore, matrix factorization and correlation-based methods designed for unpaired data require a separate feature selection step prior to integration for dimension reduction, and the method’s performance is sensitive to which genes are selected. Most existing neural network methods for multi-omics integration are based on autoencoders which, with the exception of few [25], require paired data. In general, unsupervised training of multiple autoencoders simultaneously can be very challenging without pairing information across different modalities, with finding a common embedding manifold becomes more challenging as the complexity of the data increases. Hence, current methods are limited in their ability to handle the complexity and scale that characterize multi-omics atlas data.

Here, we present a novel scalable transfer learning method, scJoint, that effectively integrates atlas-scale scRNA-seq and scATAC-seq data using a neural network approach (Figure 1a). We achieve this by taking advantage of the increasing amount of scRNA-seq data with high quality annotations and incorporating the cell type label information into a semi-supervised paradigm to train unlabeled scATAC-seq. scJoint is able to meet the challenges in integrating multi-omics atlas data through the use of (1) a *novel loss function* to explicitly incorporate dimension reduction as part of the feature engineering process in transfer learning, allowing the low dimensional features to be updated throughout training and *removing the need* for selecting highly variable genes; (2) a similarity loss that adds flexibility to the alignment of modalities when their cell types do not fully overlap; (3) weight sharing across encoders for different modalities resulting in stable training.

**Figure 1:**
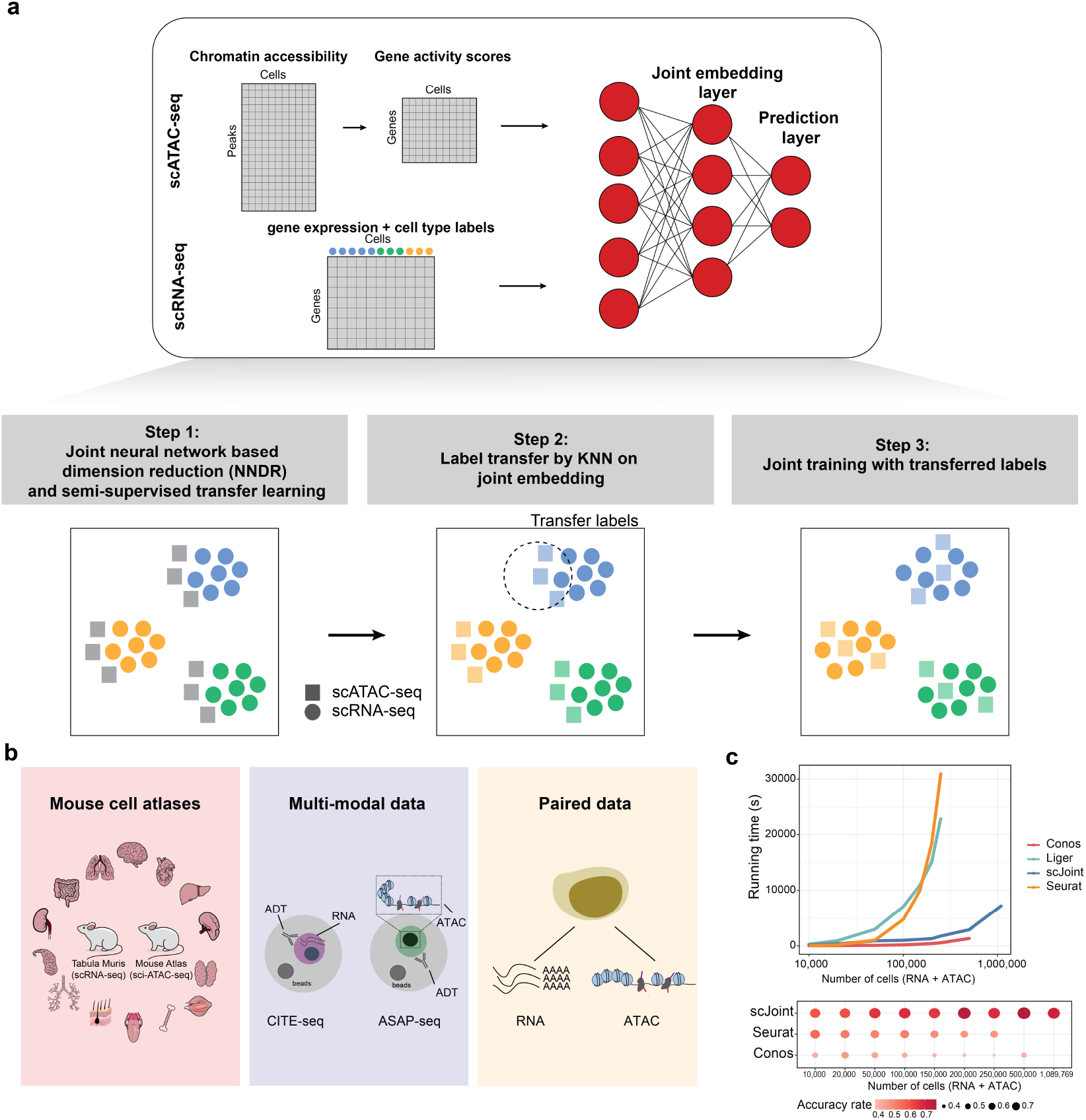
(a) Overview of scJoint. The input of scJoint consists of one (or multiple) gene activity score matrix, calculated from the accessibility peak matrix of scATAC-seq, and one (or multiple) gene expression matrix including cell type labels from scRNA-seq experiments. The method has three main steps: (1) Joint NNDR and semi-supervised transfer learning; (2) Cell type label transfer by k-nearest neighbor in joint embedding space; (3) Joint training with transferred labels. Three data collections analyzed in detail in this study: (1) Mouse cell atlases; (2) Multi-modal data from PBMC; (3) Paired data from adult mouse cerebral cortex data generated by SNARE-seq. (c) Computation time required by different methods to integrate scRNA-seq and scATAC-seq (top) and their label transfer accuracy (bottom, computed for methods with label transfer functionality). The benchmark datasets were subsampled from 54 cell types in the human fetal atlases [27, 28], where the total number of RNA and ATAC cells ranges from 10000 to 1089769. Seurat and Liger were terminated for out of memory error on datasets with 500,000 cells and more, and Conos was terminated on the 1 million cell dataset.

We illustrate scJoint’s performance in terms of label transfer accuracy, quality of joint visualizations, scalability and capacity to generalize. In particular, we highlight the scalability of scJoint through the integration of two mouse atlases [5, 26] and two human fetal atlases [27, 28]. In the latter case, scJoint required only two hours to integrate over a million cells (Figure 1c) while consistently maintaining high accuracy rates. The generalizability of scJoint to other types of single-cell data is demonstrated through a multi-modal data with paired protein measurements (CITE-seq and ASAP-seq, Figure 1b).

## Results

### scJoint for co-training labeled and unlabeled data

The core of scJoint is a semi-supervised approach to co-train labeled data (scRNA-seq) and unlabeled data (scATAC-seq), where we address the main challenge of aligning these two distinct data modalities via a common lower dimensional space. scJoint consists of three main steps (Figure 1a). Step 1 performs joint dimension reduction and modality alignment in a common embedding space through a novel neural network based dimension reduction (NNDR) loss and a cosine similarity loss respectively. The NNDR loss extracts orthogonal features with maximal variability in a vein similar to PCA, while the cosine similarity loss encourages the neural network to find projections into the embedding space so that majority parts of the two modalities can be aligned. The embedding of scRNA-seq is further guided by a cell type classification loss, forming the semi-supervised part. In Step 2, treating each cell in scATAC-seq data as a query, we identify the k-nearest neighbors (KNN) among scRNA-seq cells by measuring their distances in the common embedding space, and transfer the cell type labels from scRNA-seq to scATAC-seq via majority vote. In Step 3, we further improve the mixing between the two modalities by utilising the transferred labels in a metric learning loss. Joint visualization of the datasets is obtained from the final embedding layer using standard tools including tSNE [29] and UMAP [30]. scJoint requires simple data preprocessing with the input dimension equal to the number of genes in the given datasets after appropriate filtering. Chromatin accessibility in scATAC-seq data is first converted to gene activity scores [31, 32] allowing for the use of a single encoder with weight sharing for both RNA and ATAC.

We next compared scJoint with methods recently developed and applied to the integration of scRNA-seq and scATAC-seq, including Seurat v3 [24], Conos [23] for label transfer accuracy, and additionally Liger [21] (as a representative matrix factorization method) for evaluating the joint embedding of the two modalities.

### scJoint shows accurate and robust performance on large atlas data

We demonstrate the performance of scJoint in a complex scenario, where the heterogeneity of cell types and tissues in atlas data poses significant challenges to data integration. We applied our method to integrate two mouse cell atlases: the Tabula Muris atlas [5] for scRNA-seq data and the atlas in [26] for scATAC-seq data, containing 73 cell types (96,404 cells from 20 organs, two protocols) and 29 cell types (81,173 cells from 13 tissues) respectively (the latter including a group annotated as “unknown”), of which 19 cell types are common. We focus our initial evaluation on the subset of the atlas data containing 101,692 cells from the 19 overlapping cell types only. Here, we transferred cell type labels from scRNA-seq to scATAC-seq and compared the results with the original labels in [26] for accuracy; these original labels were also used to evaluate the quality of joint visualizations. An inspection of the tSNE plots shows our method effectively mixes the three protocols (FACS, droplet, ATAC) while providing a better grouping of the cells in terms of previously defined cell types than the other methods (Figure 2a, Supplementary Figure S1). This observation is confirmed by the quantitative evaluation metrics, with scJoint showing significantly higher cell type silhouette coefficients than all the other methods and similar modality silhouette coefficients as Seurat and Liger. Overall, scJoint has the highest median F1-score of silhouette coefficients, achieving a better trade-off between removing the technological variations in modalities and maintaining the cell type signals (Figure 2b, Supplementary Figure S2). In terms of label transfer accuracy, scJoint assigned 84% of the cells to the correct type, 14% and 13% higher than Seurat and Conos (Figure 2d, Supplementary Figure S3).

**Figure 2:**
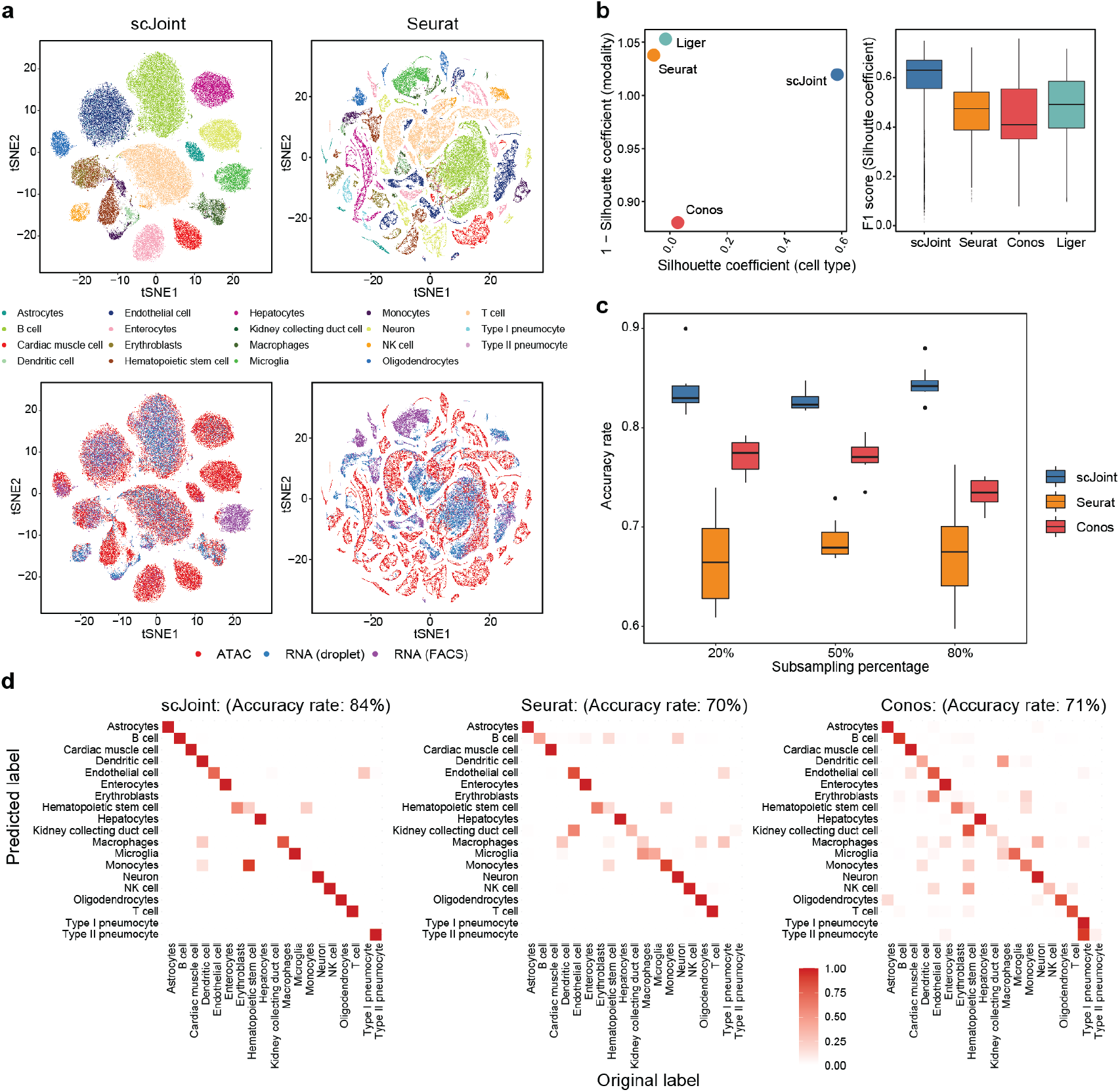
Analysis of mouse cell atlas subset data containing 19 overlapping cell types from RNA and ATAC. (a) tSNE visualization of scJoint (left column) and Seurat (right column), colored by cell types defined in [26] (first row) and three protocols (second row). (b) Scatter plot of mean silhouette coefficients for scJoint, Liger, Seurat, and Conos (left panel), where the x-axis shows the mean cell type silhouette coefficients and the y-axis shows ‘1 - mean modality silhouette coefficients’; ideal outcomes would lie in the top right corner. Boxplots of F1 scores of silhouette coefficients for scJoint, Liger, Seurat, and Conos (right panel). (c) Accuracy rates of scJoint, Seurat and Conos using 20%, 50% and 80% of cells from scRNA-seq data as training data. 10 random subsamplings were performed for each setting to generate the variance. (d) Predicted cell types and their fractions of agreement with the original cell types given in [26] for scJoint (left panel), Seurat (middle panel) and Conos (right panel). Clearer diagonal structure indicates better agreement.

To assess the robustness of the label transfer results, we performed a stability analysis on this subset of atlas data by subsampling 80%, 50%, 20% of the cells from scRNA-seq as the training data. Even when only 20% of the cells were used for training, scJoint maintained a high accuracy and small variance (Figure 2c), suggesting that scJoint is potentially applicable to situations where only a subset of the scRNA-seq data is annotated.

To evaluate scJoint’s computational efficiency on atlas-sized data, we further considered two human fetal atlases [27, 28] and created benchmark datasets by subsampling from 15 organs with 54 cell types common between scRNA-seq and scATAC-seq. The size of the datasets ranged from 10,000 to 1,089,769 cells. scJoint was significantly faster than Seurat and Liger, being the only method capable of handling over 1 million cells (2 hours using a single thread for PyTorch, Figure 1c, Supplementary Figure S4). scJoint consistently achieved much higher accuracy than the other methods, with an average 20% improvement for 100,000 or more cells (Figure 1d). Together, these results illustrate that scJoint scales well to large atlas data both in terms of computational efficiency and quality of results.

### Label transfer using highly heterogeneous atlas data refines cell type annotations in scATAC-seq

We next performed the more challenging task of integrating full atlas data, using the mouse atlases as an example. Since the scRNA-seq atlas contains more cell types than the scATAC-seq atlas, we use this application to illustrate how transferred labels can refine and provide new annotations to ATAC cells. To compare with the original labels, tSNE plots were constructed in the same way as [26], using singular value decomposition of the term frequency-inverse document frequency (TF-IDF) transformation of scATAC-seq peak matrix (Figure 3a). We observe that scJoint labels cells close together in this ATAC visualization space in a more consistent way than the other methods. Qualitatively this is supported by scJoint’s higher overall accuracy rate (77% compared with 60% for Seurat and 55% for Conos).

**Figure 3:**
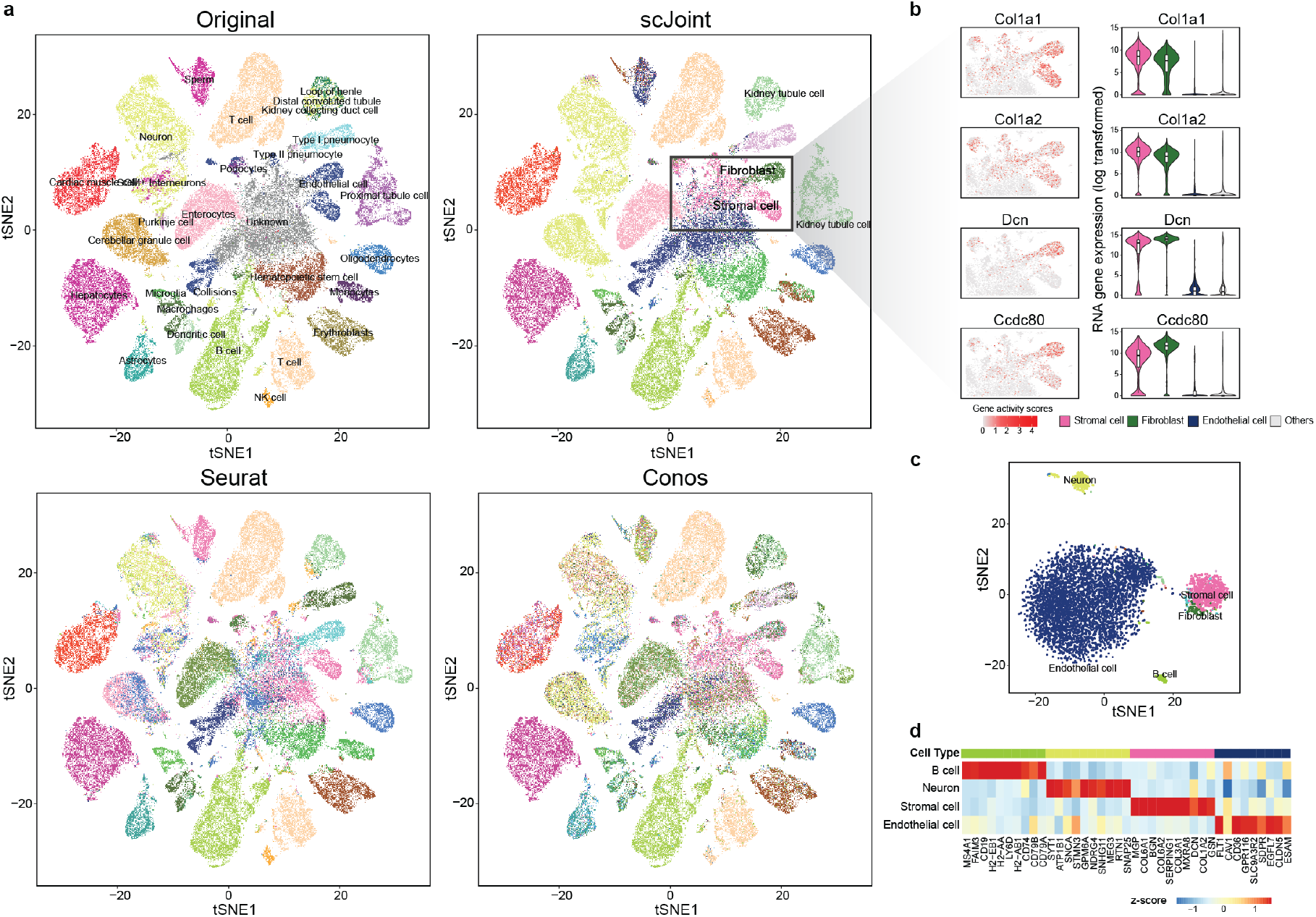
Analysis of mouse cell atlas full data. (a) A 2 *×* 2 panel of tSNE plots generated from top 100 dimensions of singular value decomposition of the TF-IDF transformed ATAC-seq data, colored by the original labels (top left), scJoint transferred labels (top right), Seurat transferred labels (bottom left), and Conos transferred labels (bottom right). (b) Marker expressions in stromal cells and fibroblasts: Col1a1, Col1a2, Dcn and Ccdc80. The left column shows the gene activity scores of the markers in ATAC-seq data (4352 stromal cells, and 1602 fibroblasts). The right column shows the log-transformed gene expression of the markers in stromal cells, fibroblasts, endothelial cells versus others; all cells here are taken from the FACS scRNA-seq data. (c) tSNE plot of cells originally labeled as ‘unknown’ and annotated by scJoint with probability scores greater than 0.80, colored by predicted cell types (5931 cells). (d) Heatmap of z-scores of average gene activity scores, calculated from cells aggregated by predicted cell types in ATAC. The rows indicate the top four predicted cell types by size. The columns indicate the top differential expressed genes of the corresponding cell type in RNA.

Examining the transferred labels further, we find scJoint labels a group of cells (originally labeled as “unknown” or “endothelials”) as “stromal cells” (4352 cells) and “fibroblasts” (1602 cells), which are two cell types not present in the original ATAC labels. These cells show high gene activity scores for Col1a1, Col1a2, Dcn and Ccdc80, all of which are markers with high expression levels in stromal cells and fibroblasts but low expression levels in endothelial cells from the scRNA-seq data (Figure 3b). Hence, the new annotations are more consistent with the marker expression levels.

More interestingly, we note scJoint allows us to annotate 5931 cells labeled as ‘unknown’ in [26] with probability score greater than 0.80. These cells are clearly clustered into groups in the tSNE visualization of scJoint’s embedding space (Figure 3c), with the main groups being endothelial cells, stromal cells, neurons and B cells. Using cell type markers identified from the scRNA-seq data, the aggregated gene activity scores of these ATAC cells show clear differential expression patterns (Figure 3d).

### scJoint enables accurate integration of single-cell multi-modal data across biological conditions

We demonstrate scJoint is capable of incorporating additional modality information to RNA-seq and ATAC-seq and applicable to experiments with different underlying biological conditions. We consider multi-modal measurements profiling gene expression levels or chromatin accessibility simultaneously with surface protein levels, which can be obtained via CITE-seq [33] and ASAP-seq [34]. We analyzed CITE-seq and ASAP-seq data from a T cell stimulation experiment in [34], which sequenced cells with these two technologies in parallel. A total of 18,088 cells were studied under two conditions: one with stimulation of anti-CD3/CD28 in the presence of IL-2 for 16 hours and the other without stimulation as control. We first clustered and annotated these cells using CiteFuse [35]. Compared to the cell type labels in the original study, we were able to identify cellular subtypes with CiteFuse, further annotating five subgroups in T cells. Next, we performed integration analysis of CITE-seq and ASAP-seq by concatenating gene expression or gene activity vectors with protein measurements. The analysis was performed in two scenarios: within the stimulated and control condition separately and across the two conditions.

In both scenarios, scJoint generated a better joint visualization of the two technologies (Figure 4a, Supplementary Figures S5, S6). In particular, in the case where stimulated and control cells are combined, subtypes of T cells (e.g. naive CD8+, effector CD8+, naive CD4+, and effector CD4+) are clearly separated while cells from the two technologies are well mixed (Figure 4a-b). The median cell type silhouette coefficient of scJoint is 0.51, outperforming the other three methods by a large margin (Seurat 0.11, Conos 0.13, and Liger -0.06). With the highest silhouette coefficient F1 scores (median F1 score: 0.59) representing a 16% - 28% improvement over the other methods, scJoint demonstrates the best balance between removing technical variations and preserving biological signals (Figure 4c, Supplementary Figure S7).

**Figure 4:**
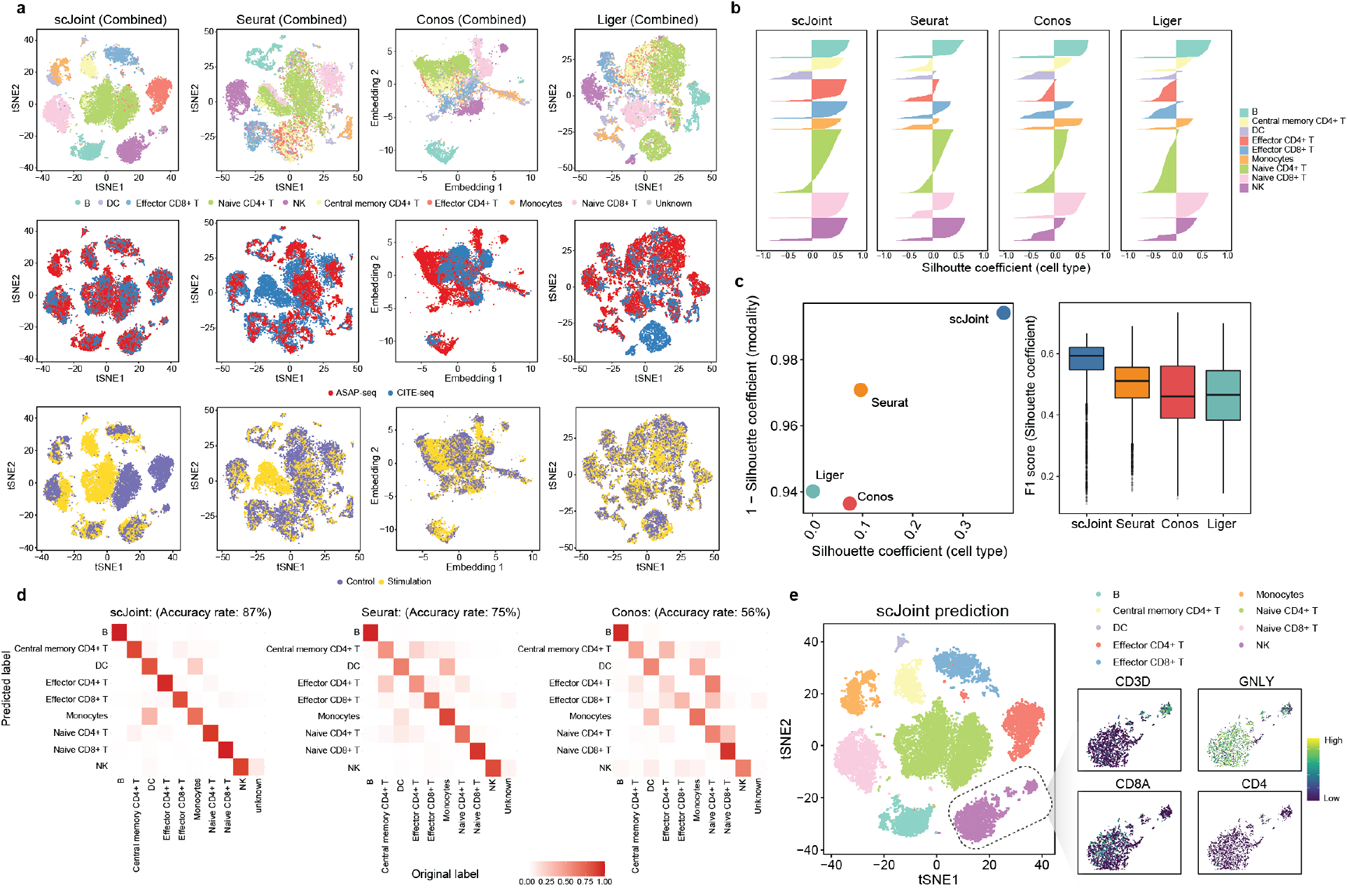
Integration of multi-modal PBMC data across biological conditions. (a) tSNE visualization of scJoint (first column), Seurat (second column), Conos (third column) and Liger (fourth column) of PBMC data generated from CITE-seq and ASAP-seq, colored by cell type obtained from CiteFuse and manual annotations (first row), technology (second row), and biological condition (third row). (b) Barplots of cell type silhouette coefficients for scJoint, Seurat, Conos and Liger for all cells, colored by cell type. Larger values on the x-axis indicate better grouping. Scatter plot of mean silhouette coefficients for scJoint, Seurat, Conos and Liger (left), where the x-axis denotes the mean cell type silhouette coefficients, and the y-axis denotes 1 - mean modality silhouette coefficients; ideal outcomes would lie in the top right corner. Boxplots of F1 scores of silhouette coefficients for scJoint, Liger, Seurat, and Conos (right). (d) Heatmaps comparing the original labels and the transferred labels of scJoint, Seurat and Conos. Clearer diagonal structure indicates better agreement. (e) tSNE visualization of scJoint colored by the predicted cell types with gene expression levels of CD3D, NKG7, CD8A and CD4 in natural killer cells.

Moreover, scJoint achieves higher accuracy in label transfer under all scenarios (88% in control, 84% in stimulation, and 87% in the combined case), compared with Seurat (80% in control, 79% in stimulation, and 75% combined) and Conos (53% in control, 67% in stimulation, and 56% in combined) (Figure 4d and Supplementary Figure S8). In addition, the transferred labels of scJoint from the two scenarios (control / stimulation alone, and combined) are highly consistent, with 95% of cells having the same annotation, substantially greater than Seurat (84%) and Conos (59%) (Supplementary Figure S9).

### Integration of multi-modal data with scJoint captures additional biological signals in cell types and conditions

In the combined analysis of stimulation and control, we find that the joint embedding generated by scJoint contains additional information that allows for the identification of a cellular subtype. In the CiteFuse annotation of ASAP-seq data, we labeled one cluster of 142 cells with ambiguous marker expression as “unknown”. Interestingly, in the joint visualization of scJoint, while these “unknown” cells are labeled as “natural killer cells (NK)” by label transfer, they are still clearly separated from the majority of NK cells and form a small cluster together with cells from CITE-seq. We then examined the gene and protein expression levels of NK cell and T cell markers in this subgroup. We find these cells have high expression of CD3 and GNLY at gene level as well as CD3, CD56, CD57, and CD244 at protein level, but low expression of CD8A and CD4. This suggests these cells may be natural killer T cells, a minority of immune cells in PBMC sample (Figure 4e, Supplementary Figure S10) [36]. By contrast, although these cells lack CD8 expression, the other methods are unable to distinguish them from effector CD8+ T cells in their visualizations (Figure 4e, Supplementary Figure S11).

Lastly, by appropriately aligning the two technologies in the embedding space, scJoint is able to reveal the biological difference between stimulation and control within the same cell type. In the joint visualization of scJoint, three subtypes of T cells (naive CD4+, naive CD8+, effector CD4+) are less well mixed between the two conditions than the other cell types, consistent with the stimulation experiment aiming to activate T cells. In particular, the naive CD4+ T cells show the most notable separation between the two conditions (Figure 4a). We then performed differential expression analysis of the scRNA-seq part of CITE-seq within each cell type across the two conditions using MAST [37]. We find that the naive CD4+ T cells have the largest number of unique differentially expressed genes (FDR *<* 0.01) (Supplementary Figure S12a). Similarly, differential proteins analysis of both CITE-seq and ATAC-seq using wilcoxon rank sum test on the log-transformed protein abundances also suggests that naive CD4+ T cells have the most unique differential proteins compared with other cell types (FDR *<* 0.01) (Supplementary Figure S12b-c).

### scJoint shows versatile performance on paired measurements of scRNA-seq and scATAC-seq

Although scJoint is designed for integrating unpaired data, it is still directly applicable to paired data. Such an application also enables us to compare its performance with methods that incorporate pairing information and use the pairing information to validate the label transfer results. We consider the integration of adult mouse cerebral cortex data generated by SNARE-seq [14], a technology that can profile gene expression and chromatin accessibility in the same cell. In addition to Seurat and Liger, we compared scJoint with two other methods designed specifically for paired data, scAI [16] and MOFA+ [17]. In our assessment, all the unpaired methods (scJoint, Seurat, Liger) treat the RNA and ATAC parts of SNARE-seq as two separate datasets, while the paired methods take the pairing information into account. We find that scJoint is able to provide clear groupings of cells according to cellular subtypes (Figure 5a) and achieves comparable or better cell type silhouette coefficients (Figure 5b) than the paired methods. This suggests that scJoint is versatile enough to be applied to paired data, which are becoming increasingly popular.

**Figure 5:**
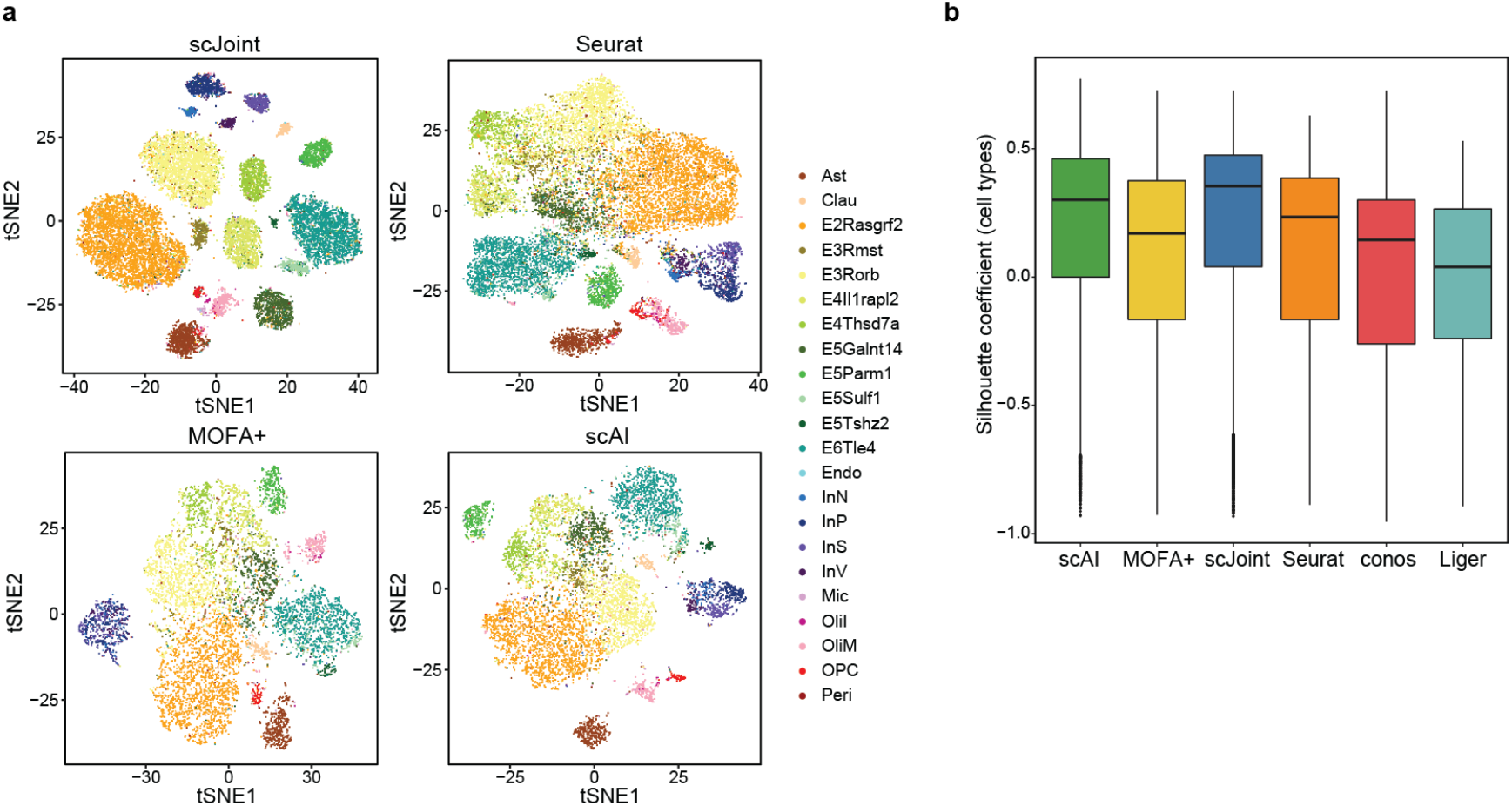
Analysis of paired gene expression and chromatin accessibility data from SNARE-seq. (a) tSNE visualization of SNARE-seq data for scJoint, Seurat, MOFA+ and scAI, colored by cell types given in [14]. All unpaired methods treat the RNA and ATAC parts of SNARE-seq as two separate data. (b) Boxplots of cell type silhouette coefficients for scJoint, Seurat, Conos and Liger, colored by methods.

Comparing the performance among the unpaired methods, scJoint has the highest medians in cell type silhouette coefficients and F1-scores (Figure 5b, Supplementary Figure S14). For label transfer, scJoint achieves an accuracy rate of 70.9%, retaining better performance than the other two methods (70.1% for Seurat and 49.5% for Conos). Looking closer at the performance in each cell type, scJoint performs the best in 10 out of 22 cell types in terms of F1 scores for classification (Supplementary Figure S15). Together, these results suggest that scJoint performs the best among the unpaired methods and on par with the paired methods, despite treating paired data as separate.

## Discussion

scJoint approaches the integration of scRNA-seq and scATAC-seq as a domain adaptation problem in transfer learning, using the same neural network to co-train labeled data from the source domain (RNA) and unlabeled data from the target domain (ATAC) following a different distribution. scRNA-seq data serve as a natural source domain for transferring information to other modalities due to rapidly growing collections of annotated public data and RNA-focused computational tools that can output accurate classifications [38]. Using multiple cell atlases and multi-modal data with protein measurements, we demonstrate scJoint achieves significantly higher label transfer accuracy and provides better joint visualizations than other methods even when 1) the data is highly complex and heterogeneous and 2) meaningful biological conditions are mixed with technical variations. We have shown that integrative analysis of single-cell multi-omics data by scJoint facilitates re-annotation of cell types in scATAC-seq and discovery of new subtypes not present in training data.

scJoint provides a concise training framework with one main tuning parameter in the construction of cosine similarity loss. As shown in Supplementary Figure S16a, our results are quite stable with respect to the choice of this parameter. Similar to other methods based on neural networks, the number of hidden nodes in the architecture and other optimization details can be considered tunable as well, although they do not appear to affect our results (Supplementary Figure S16b).

The superior performance and robustness of scJoint illustrate its utility as a tool to automatically label cells from other modalities given an annotated scRNA-seq database. By embedding all cells in a common lower dimensional space, scJoint assigns a probability score to a cell type prediction by combining the softmax probabilities of its nearest neighbors. As we vary the level of cutoff, the accuracy of scJoint still consistently outperforms the other methods (Supplementary Figure S17). The robustness of scJoint was demonstrated through subsampling experiments, where the stability of our results implies the method can be applied to partially labeled databases. Despite being a semi-supervised method guided by labeled data, the dimension reduction component in our design lends it sufficient flexibility to preserve implicit data signals, including biological variations induced by experimental conditions and additional cellular subtypes. One can conceivably extend scJoint to an unsupervised setting, replacing the softmax prediction layer with a decoder minimizing reconstruction loss.

Although designed for unpaired data, scJoint is still directly applicable to paired data and generates joint visualizations with cells coherently grouped by cell types. In the current training scheme, the pairing information between RNA and ATAC is only used to validate the label transfer results. We expect that adapting scJoint to take paired vectors during training would enhance its performance on this type of data, and this would be especially useful in the unsupervised setting mentioned above.

We have focused on scATAC-seq as an example of epigenomic data, but in principle scJoint extends to other modalities such as methylation data, provided the input can be summarized as gene-level scores. While the gene-level summaries are amenable to generalization and widely adopted by unpaired integration methods, this step itself is also a limitation as improper aggregation can incur information loss. Extending scJoint to directly handle epigenomic data at locus level will require designing a separate encoder that is suitable for the high dimensionality and remains easy to train, and we will pursue this for future work.

In summary, we have developed scJoint as a generalizable transfer learning method for performing integrative analysis of atlas-scale single-cell multi-omics data. scJoint was shown to effectively integrate multiple types of measurements from both unpaired or paired profiling, outperforming other methods in label transfer accuracy and providing joint visualizations that remove technical variations while preserving meaningful biological signals. scJoint’s ability to integrate multi-omics data by capturing various aspects of cell characteristics unique to different data modalities will facilitate a more comprehensive view of cell functions and cell communications.

## Methods

### Architecture and training of scJoint

The neural network in scJoint consists of one input layer and two fully connected layers. The input layer has dimension equal to the number of genes common to the expression matrix of scRNA-seq and the gene activity matrix of scATAC-seq, after simple filtering (see Data preprocessing). Now that the two modalities have matching input features, we co-train them using the same encoder which is equivalent to weight sharing. The first fully connected layer has 64 neurons with linear activation and serves as the joint low dimensional embedding space that captures aligned features from all cells. visualizations of clustering structure can be obtained by applying tSNE or UMAP to the output of the embedding layer. The second fully connected layer has dimension equal to the number of cell types in scRNA-seq data. Through a softmax transformation, this layer outputs a probability vector for cell type prediction. For cells in scRNA-seq, this layer can be trained in a supervised fashion using the cross entropy loss.

Given *S* scRNA-seq experiments with expression matrices and *T* scATAC-seq experiments with gene activity score matrices, with *S* and *T* representing the number of different batches whose technical variations need to be removed. Assume suitable intersections have been taken so that all matrices have the same set of genes. Let 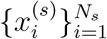 be the expression profiles of cells after preprocessing from a scRNA-seq dataset indexed by *s* ∈ *{*1,. .., *S}*, and 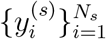 be the corresponding cell type annotations. Here each 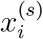 is a *G*-dimensional vector, where *G* is the number of genes; 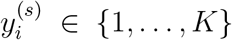, where *K* is the number of cell types; *N*_*s*_ is the number of cells in experiment *s*. Similarly, let 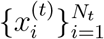 be the vectors of gene activity scores after preprocessing from the *t*-th scATAC-seq dataset with *N*_*t*_ cells (*t* ∈ *{*1,. .., *T}*), whose cell types are unlabeled. The neural network is parametrized by a set of weights and biases, collectively denoted *θ*. Let 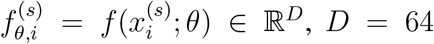, be the output of the embedding layer when the input 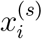 has gone through a transformation of *f* parametrized by *θ*. Similarly 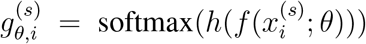, where *h* denotes the output from the prediction layer that goes through the softmax transformation. Thus 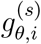 is a probability vector after the softmax transformation. 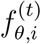 and 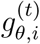 are defined in the same way for input 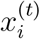 from scATAC-seq.

The training of scJoint consists of three steps.

### Step 1: Joint neural network based dimension reduction (NNDR) and semi-supervised transfer learning

We first perform joint dimension reduction and feature alignment by imposing suitable loss functions on the outputs of the two fully connected layers. A mini-batch *B*_0_ of data for training is constructed by sampling equal-sized subsets of cells from each dataset, that is, 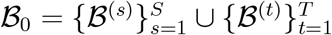, where each subset *ℬ*^(*s*)^ (or *ℬ*^(*t*)^) has *B* cells.

1. *NNDR Loss*. In a spirit similar to PCA, the NNDR loss aims to capture low dimensional, orthogonal features when projecting each data batch into the embedding space. For now we omit the dataset-specific superscript with the understanding that this loss function is applied to each *ℬ*^(*s*)^ and *ℬ*^(*t*)^. Given input vectors *{x*_*b*_*}*_*b*∈*ℬ*_, define 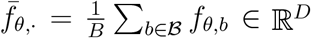, and ∑ _*θ*,·_ as the sample correlation matrix. The NNDR loss is:

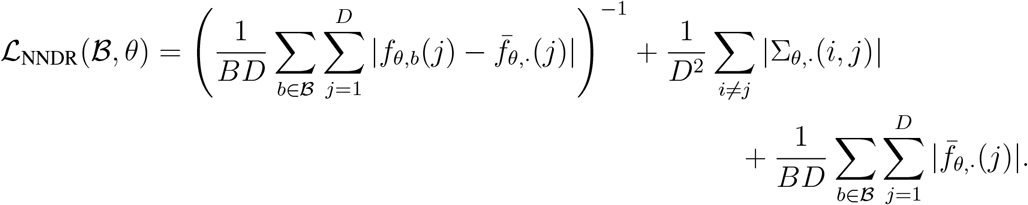 Note that to minimize this loss, we maximize the variability within each coordinate (inverse of the first term) and minimize the correlation between all coordinate pairs (the second term) to achieve orthogonality. The last term tries to fix the means of all coordinates near zero for model identifiability, preventing *θ* from drifting to unstable regions of the parameter space.
2. *Cosine similarity loss*. This loss is applied to the embedding layer outputs from *ℬ*^(*t*)^ and 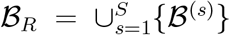, for every *t*, and attempts to maximize the similarity between best aligned ATAC and RNA data pairs. Let *p* be the fraction of data pairs we expect to have high cosine similarity scores. Setting *p<* 1 accounts for situations where RNA and ATAC do not share all their cell types. We set *p* = 0.8 for all the results presented in the paper, and our results appear to be stable with respect to this parameter (Supplementary Figure S16a) when the cell types fully overlap. Recall that for a pair of general vectors (*u, v*), the cosine similarity is defined as cos(*u, v*)= ⟨*u, v*⟩*/*(∥*u*∥∥*v*∥). For each 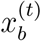 with *b* ∈ *ℬ*^(*t*)^, we find the corresponding *i*(*b*) ∈ *ℬ*_*R*_ with input *x*_*i*(*b*)_ that maximizes 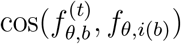. From *ℬ*^(*t*)^, we then choose the top *p* fraction of cells with the highest cosine score and denote the index set *I*_*p*_. (*I*_*p*_ has size ⌊*Bp*⌋.) The loss is given by

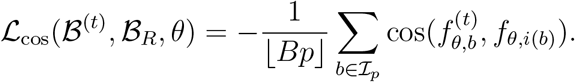
3. *Cross entropy loss*. For every *B*^(*s*)^ with cell type annotations 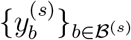, we apply the cross entropy loss to the prediction layer after softmax transformation to supervise the learning of scRNA-seq datasets:

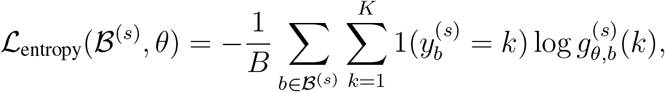

where 1(·) is an indicator function.

In Step 1, the final loss function we minimise with respect to *θ* for a mini-batch *ℬ*_0_ is

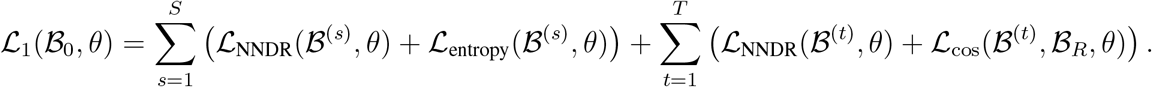

### Step 2: Cell type label transfer by KNN in joint embedding space

The output of Step 1 is a joint embedding space that has roughly aligned RNA and ATAC with cells from either modality lying close if they have similar low dimensional representations in this space. Therefore using the embedding vectors for cells in all the datasets and calculating the Euclidean distances, we can determine the KNN among all RNA cells for each cell *i* in ATAC; denote this set of RNA cells 𝒩 (*i*). The cell type label of *i* is estimated via majority vote using *{y*_*j*_*}*_*j*∈𝒩 (*i*)_. All the results in the paper were obtained from using 30 nearest neighbors. Let the majority cell type be *k*^*\**^, then the probability score of cell type prediction for cell *i* in ATAC is an average of its nearest neighbors in RNA. Since for each 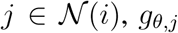 is already a probability vector after the softmax transformation, we take 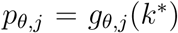 as the probability score of RNA cell *j* in the majority class ℳ(*i*) ⊂ 𝒩 (*i*). For other *j* ∈ 𝒩 (*i*)*\*ℳ(*i*), we threshold the probability score as 0. Then the probability score of ATAC cell *i* is calculated as

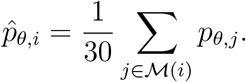

### Step 3: Joint training with transferred cell type labels

In the final step of the training, we refine the joint embedding space and improve mixing of cells from the same cell type using the transferred labels from Step 2. We include an additional loss function commonly used in metric learning for enhancing embedded clustering structure given labeled data. The other loss functions and network architecture remain the same as Step 1 with ATAC cells and their transferred labels added to *ℒ*_entropy_.

For each cell type *k* ∈ *{*1,. .., *K}*, we initialize the class center *c*_*k*_ ∈ ℝ^*D*^ randomly. We construct mini-batches of cells from all the datasets in the same way as Step 1. Now that all cells have cell type labels (given or transferred), for convenience we will refer to cells in a mini-batch *ℬ*_0_ without explicitly labeling which dataset they come from. For a given *ℬ*_0_, we first update the class centers by taking the average of *c*_*k*_ and *{f*_*θ,b*_*}* with *b* ∈ *ℬ*_0_ and *y*_*b*_ = *k*. Let the updated centers be 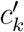. As the number of mini-batches grows, the influence of the initial *c*_*k*_ becomes negligible. The metric learning loss we use is the center loss:

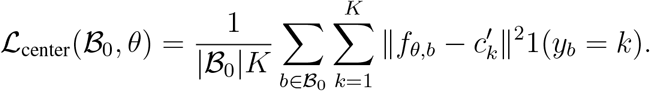

The total loss function we minimise in Step 3 is given by

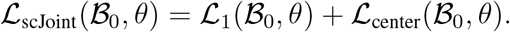

We perform a final round of majority vote by KNN using distances in the embedding space. If the prediction of any ATAC cell is different from Step 2, we update both its prediction and probability score in the same way as Step 2. Before visualization with tSNE, all embedding vectors are normalized using *L*_2_ norm.

### Training details

The batch size *B* was set to 256 in all cases. The other training details including learning rate and number of training epochs used in each dataset can be found in Table S1. We started all the training with learning rate set to 0.01, since a large learning rate has the benefit of faster training. However, if the values of the loss functions were observed to have too much fluctuation, we decreased the learning rate to 0.001 for more stable training.

### Data preprocessing

- *Mouse atlas data*. The processed gene expression matrix and the cell type annotation of the Tabula Muris mouse data of scRNA-seq were downloaded from https://tabula-muris.ds.czbiohub.org/, which have 41965 cells from protocol fluorescence-activated cell sorting (FACS) and 54439 cells from microfluidic droplets (droplet). The quantitative gene activity score matrix and the cell type annotation of Mouse sci-ATAC-seq Atlas were downloaded from https://atlas.gs.washington.edu/mouse-atac/, including 81173 cells in total. The number of common genes between two modalities is 15519. We manually checked the cell type annotations from the original studies and re-annotated the labels such that the naming convention is consistent across the datasets. For example, the cell type “Cardiac muscle cell” in the sci-ATAC-seq dataset was changed to “Cardiomyocytes”. We also combined some of the cellular subtypes in the sci-ATAC-seq data to increase the percentage of overlapping labels between two atlases for evaluation. More specifically, we combined “Regulatory T cell” and “T cell” into “T cell”; “Immature B cell”, “Activated B cell” and “B cell” into “B cell”; “Excitatory neurons” and “Inhibitory neurons” into “Neuron”.
- *Human fetal atlas data*. The scRNA-seq data of the human fetal atlas data was downloaded from GSE156793, including both raw gene expression and cell type information [27]. The scATAC-seq data was downloaded from GSE149683, and the gene activity matrices were extracted from the Seurat objects provided [28]. There are 54 cell types common between the two human fetal atlases. In our computational benchmarking analysis, we only included cells from the common cell types, resulting in a total of 656,074 cells from the scATAC-seq data. To construct a balanced scRNA-seq training set, for cell type *i* with number of cells *n*_*i*_ *>* 10, 000, we subsampled max*{*0.05*n*_*i*_, 10, 000*}* cells; all cells were included for cell types with less than 10,000 cells. This resulted in 433,695 cells from the scRNA-seq data.
- *SNARE-seq data*. The SNARE-seq data from adult mouse cerebral cortex was downloaded from the National Center for Biotechnology Information (NCBI) Gene Expression Omnibus (GEO) accession number GSE126074 [14], with both raw gene expression and DNA accessibility measurements available for the same cell. The fastq files were downloaded from the Sequence Read Archive (SRA) for SRP183521. We first derived the fragment files from the fastq files using `sinto fragments` (sinto v0.7.2), and then generated the gene activity matrix using Signac (v1.1.0.9000) [32]. The cell type information was obtained from the original study [14]. We filtered out the cells that were originally labeled as “Misc” (cells of miscellaneous cluster), resulting in a dataset with 9190 cells and 15725 genes for the integrative analysis.
- *Multi-modal data (CITE-seq and ASAP-seq PBMC data)*. The ASAP-seq and CITE-seq data were downloaded from GEO accession number GSE156478 [34], which included the fragment files and antibody-derived tags (ADTs) matrices for ASAP-seq, the raw unique molecular identifier (UMI) and ADT matrices for CITE-seq, from both control and stimulated conditions. The gene activity matrices for ASAP-seq were generated by Signac. Most of the thresholds we used for quality control metrics were consistent with those in the original paper [34]. The control and stimulated CITE-seq were filtered based on the following criteria: mitochondrial reads greater than 10%; number of expressed genes less than 500; total number of UMI less than 1000; total number of ADTs from the rat isotype control greater 55 and 65 in the control and stimulated conditions respectively; total number of UMI greater than 12,000 and 20,000 for the control and stimulated conditions respectively; total number of ADTs less than 10,000 and 30,000 for control and stimulated conditions respectively. We further filtered out cells that were classified as doublets in original study. For the ASAP-seq data, we filtered out cells with the number ADTs more than 10,000 and number of peaks more than 100,000. Finally, 4502 cells (control) and 5468 cells (stimulated) from ASAP-seq, 4644 cells (control) and 3474 cells (stimulated) from CITE-seq were included in the downstream analysis. The number of common genes across the four matrices is 17441 and the number of common ADTs is 227. We used CiteFuse to integrate the peak matrix or gene expression matrix with their corresponding protein expression and obtain clustering for ASAP-seq and CITE-seq within each condition separately [35]. For ASAP-seq, the similarity matrices of the chromatin accessibility are calculated by applying the Pearson correlation to the TF-IDF transformation of the peak matrix. We then followed the procedure described in [39] to annotate the clusters.

For scJoint, all the gene expression matrices and gene activity score matrices were binarized as 0 or 1, with 1 representing any non-zero original values, as the final input for training. Binarization scales the two modalities so that their distributions have the same range and reduces the noise level in the data for easier co-training.

### Settings used in other methods

For the unpaired data (mouse cell atlases and multi-modal data from CITE-seq and ASAP-seq), we benchmarked the performance of scJoint against three other methods designed for integrating unpaired single-cell multi-modal data: Seurat (v3), Conos and Liger. We compared the label transfer accuracy with Seurat and Conos and the joint visualizations with all three methods. For the paired data (SNARE-seq), we further compared joint visualizations with two methods specifically designed for paired data, scAI and MOFA+. For all the unpaired methods, we used gene activity matrices derived from the above data preprocessing step as input for scATAC-seq. For the two paired methods, we used the peak matrices of scATAC-seq data as input. Detailed settings used in each method are as follows.

- *Seurat*. R package Seurat v3.2.0 [24] was used for all the datasets. The raw count matrix of scRNA-seq and unnormalized gene activity score matrix of scATAC-seq were used as input, which were then normalized using the `NormalizeData` function in Seurat. Noted that for the CITE-seq and ASAP-seq data, the input was a concatenated matrix of log-transformed normalized gene expression data/gene activity score matrix and log-transformed ADTs matrix. Top 2000 most variable genes were selected from scRNA-seq using `FindVariableFeatures` with `vst` as method. To identify the anchors between scRNA-seq and scATAC-seq data, `FindTransferAnchors` function was used with “cca” as reduction method. The scATAC-seq data was then imputed using `TransferAnchors` function, where the anchors were weighted by latent semantic indexing (LSI) reduced dimension of scATAC-seq. Principal component analysis was then performed on the merged matrix of scRNA-seq data and imputed scATAC-seq data. For all the datasets, 30 principal components (PCs) were used for joint visualization with tSNE (function `RunTSNE`). For the mouse cell atlas data, we first integrated the two scRNA-seq datasets (FACS and droplet) using `FindIntegrationAnchors` and `IntegrateData`, and then the integrated matrix was scaled using `ScaleData` and used as reference to find transfer anchors.
- *Conos*. R package conos v1.3.1 [23] was used for all the datasets. Function `basicP2proc` in `pagoda2` package (v0.1.2) was performed to process the raw count matrix of scRNA-seq and unnormalized gene activity score matrix of scATAC-seq. The joint graph was built using `buildGraph` with `k=15`, `k.self=5`, and `k.self.weigh=0.01`, which were set as suggested in the tutorial for integrating RNA and ATAC (http://pklab.med.harvard.edu/peterk/conos/atac_rna/example.html). The joint visualization of scRNA-seq and scATAC-seq were generated using largeVis by `embedGraph`, which is the default visualization in Conos.
- *Liger*. R package liger v0.5.0 [21] was used for the datasets. The raw count matrix of scRNA-seq and unnormalized gene activity score matrix of scATAC-seq were used as input, which were normalized using `normalize` function in liger. Highly variable genes were selected using the scRNA-seq. For the mouse cell atlas data, both FACS and droplet scRNA-seq data were used to select features. For all the datasets, number of factors was set to 20 in `optimizeALS`. tSNE was then performed on the normalized cell factors to generate the joint visualization of scRNA-seq and scATAC-seq (function `runTSNE` in liger).
- *scAI*. R package scAI v1.0.0 [16] was used for the integration of SNARE-seq data. The raw count matrix of scRNA-seq and raw peak matrix of scATAC-seq were used as input. We ran scAI using `run scAI` by setting the rank of the inferred factor set as 20, `do.fast = TRUE`, and `nrun = 1`, with other parameters set as default, as suggested in the pipeline in the github repository. tSNE plots were generated using `reducedDims` function in scAI.
- *MOFA+*. R package MOFA2 v1.0 [17] was used for the integration of SNARE-seq data. Following the suggested integration tutorial for SNARE-seq in the github repository, we first selected top 2500 most variable genes using `FindVariableFeatures` in Seurat package with `vst` as method and top 5000 most variable ATAC peaks with `disp` as method. By subsetting the counts matrix of scRNA-seq and peak matrix of scATAC-seq with the selected features, we ran MOFA+ by setting the number of factors as 10, with other parameters set as default. tSNE plots were generated using `run tsne` function in MOFA2.

## Evaluation metrics

### Joint embedding evaluation - Silhouette coefficients

To evaluate whether the joint embeddings from different methods show clustering structure reflecting biological signals or technical variations, we calculated the silhouette coefficient for each cell by considering two different groupings: (1) grouping based on the modalities (scRNA-seq or scATAC-seq), called the modality silhouette coefficient (*s*_*modality*_); (2) grouping based on known cell types, called the cell type silhouette coefficient (*s*_*cellTypes*_). Note that for the atlas data, we consider FACS and droplet in scRNA-seq as two distinct technologies and the modality silhouette coefficient has three groups (FACS, droplet, ATAC) in the calculation. For SNARE-seq, the paired methods (scAI and MOFA+) have no modality silhouette coefficients since each cell has one paired profile of RNA and ATAC. An ideal joint visualization should have low modality silhouette coefficients, suggesting the removal of the technical effect, and large cell type silhouette coefficients, indicating the cells are grouped by cell types. The euclidean distance for all methods except Conos is obtained from the tSNE embedding. For Conos, the distance is obtained from the `largeVis` embedding, which is the method’s default output.

We then summarize the two silhouette coefficients by calculating an F1-score as follows:

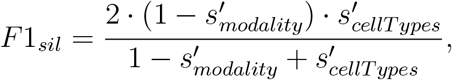

where *s*^*′*^ = (*s* + 1)*/*2. A higher F1 score indicates better performance in the alignment of the modalities as well as the preservation of biological signals.

### Accuracy evaluation of transferred labels

We evaluated the accuracy of label transfer from two aspects: (1) Overall accuracy rate; (2) Cell type classification F1-score. The overall accuracy rate was computed only accounting for the common cell types between scRNA-seq and scATAC-seq data. The cell type classification F1-score is the harmonic mean of precision and recall of each cell type.

### Running time evaluation

The running time evaluation was performed using one core and one GPU on a research server with dual Intel (R) Xeon(R) Gold 6148 Processor (40 total cores, 768 GB total memory) and dual RTX2080TI GPUs. Using the preprocessed human fetal atlas data, we created benchmarking datasets with 5000, 10000, 25000, 50000, 75000, 100000, 125000 and 250000 cells from scRNA-seq and scATAC-seq data respectively. We further ran scJoint on the whole preprocessed data with 433,695 cells from scRNA-seq and 656,074 cells from scATAC-seq. In this case, the other three methods failed to run due to an out of memory error. For each method, we measured total running time as the running time of feature selection, label transfer and joint embedding construction of scATAC-seq and scRNA-seq. The training details for scJoint are listed in Table S2. Following common practice in neural network training, we increased the batch size as the number of training data increased.

### Software availability

scJoint was implemented using PyTorch (version 1.0.0) with code available at https://github.com/SydneyBioX/scJoint.

## Acknowledgments

The authors gratefully acknowledge the following funding sources: Research Training Program Tuition Fee Offset and Stipend Scholarship and Chen Family Research Scholarship to Y.L.; Australian Research Council Discovery Project grant (DP170100654) to J.Y.H.Y.; Australian Research Council DECRA Fellowship (DE180101252) to Y.X.R.W; NIH grants R01 HG010359 and P50 HG007735 to W.H.W.

## Author contributions

T.W., W.H.W. and Y.X.R.W. conceived and designed this project; Y.L., T.W. and S.W. performed data preprocessing, model development, and evaluation of results; J.Y.H.Y., W.H.W. and Y.X.R.W. supervised the execution; Y.L., J.Y.H.Y., W.H.W. and Y.X.R.W. wrote the manuscript. All authors read and approved the manuscript.

## Conflict of interest

The authors declare that they have no conflict of interest.

## Notes

### Competing Interest Statement

The authors have declared no competing interest.

## References

1. Stuart, T. & Satija, R. Integrative single-cell analysis. Nature Reviews Genetics 20, 257–272 (2019).

2. Berger, S. L. The complex language of chromatin regulation during transcription. Nature 447, 407–412 (2007).

3. Klemm, S. L., Shipony, Z. & Greenleaf, W. J. Chromatin accessibility and the regulatory epigenome. Nature Reviews Genetics 20, 207–220 (2019).

4. Pott, S. & Lieb, J. D. Single-cell ATAC-seq: strength in numbers. Genome Biology 16, 172 (2015).

5. Schaum, N. et al. Single-cell transcriptomics of 20 mouse organs creates a Tabula Muris: The Tabula Muris Consortium. Nature 562, 367 (2018).

6. Regev, A. et al. Science forum: the human cell atlas. Elife 6, e27041 (2017).

7. Lopez, R., Regier, J., Cole, M. B., Jordan, M. I. & Yosef, N. Deep generative modeling for single-cell transcriptomics. Nature methods 15, 1053–1058 (2018).

8. Wang, J. et al. Data denoising with transfer learning in single-cell transcriptomics. Nature methods 16, 875–878 (2019).

9. Lin, Y. et al. scMerge leverages factor analysis, stable expression, and pseudoreplication to merge multiple single-cell RNA-seq datasets. Proceedings of the National Academy of Sciences 116, 9775–9784 (2019).

10. Korsunsky, I. et al. Fast, sensitive and accurate integration of single-cell data with Harmony. Nature methods, 1–8 (2019).

11. Wang, T. et al. BERMUDA: a novel deep transfer learning method for single-cell RNA sequencing batch correction reveals hidden high-resolution cellular subtypes. Genome biology 20, 1–15 (2019).

12. Amodio, M. et al. Exploring single-cell data with deep multitasking neural networks. Nature methods, 1–7 (2019).

13. Xiong, L. et al. SCALE method for single-cell ATAC-seq analysis via latent feature extraction. Nature communications 10, 1–10 (2019).

14. Chen, S., Lake, B. B. & Zhang, K. High-throughput sequencing of the transcriptome and chromatin accessibility in the same cell. Nature biotechnology 37, 1452–1457 (2019).

15. Cao, J. et al. Joint profiling of chromatin accessibility and gene expression in thousands of single cells. Science 361, 1380–1385 (2018).

16. Jin, S., Zhang, L. & Nie, Q. scAI: an unsupervised approach for the integrative analysis of parallel single-cell transcriptomic and epigenomic profiles. Genome biology 21, 1–19 (2020).

17. Argelaguet, R. et al. MOFA+: a statistical framework for comprehensive integration of multi-modal single-cell data. Genome Biology 21, 1–17 (2020).

18. Welch, J. D., Hartemink, A. J. & Prins, J. F. MATCHER: manifold alignment reveals cor- respondence between single cell transcriptome and epigenome dynamics. Genome biology 18, 1–19 (2017).

19. Amodio, M. & Krishnaswamy, S. MAGAN: Aligning biological manifolds. arXiv preprint 1803.00385 (2018).

20. Liu, J., Huang, Y., Singh, R., Vert, J.-P. & Noble, W. S. Jointly embedding multiple single- cell omics measurements. BioRxiv, 644310 (2019).

21. Welch, J. D. et al. Single-cell multi-omic integration compares and contrasts features of brain cell identity. Cell 177, 1873–1887 (2019).

22. Duren, Z. et al. Integrative analysis of single-cell genomics data by coupled nonnegative matrix factorizations. Proceedings of the National Academy of Sciences 115, 7723–7728 (2018).

23. Barkas, N. et al. Joint analysis of heterogeneous single-cell RNA-seq dataset collections. Nature methods 16, 695–698 (2019).

24. Stuart, T. et al. Comprehensive integration of single-cell data. Cell 177, 1888–1902 (2019).

25. Dai Yang, K. et al. Multi-domain translation between single-cell imaging and sequencing data using autoencoders. Nature Communications 12, 1–10 (2021).

26. Cusanovich, D. A. et al. A single-cell atlas of in vivo mammalian chromatin accessibility. Cell 174, 1309–1324 (2018).

27. Cao, J. et al. A human cell atlas of fetal gene expression. Science 370 (2020).

28. Domcke, S. et al. A human cell atlas of fetal chromatin accessibility. Science 370 (2020).

29. Maaten, L. v. d. & Hinton, G. Visualizing data using t-SNE. Journal of machine learning research 9, 2579–2605 (2008).

30. McInnes, L., Healy, J. & Melville, J. Umap: Uniform manifold approximation and projec- tion for dimension reduction. arXiv preprint 1802.03426 (2018).

31. Pliner, H. A. et al. Cicero predicts cis-regulatory DNA interactions from single-cell chro- matin accessibility data. Molecular cell 71, 858–871 (2018).

32. Stuart, T., Srivastava, A., Lareau, C. & Satija, R. Multimodal single-cell chromatin analysis with Signac. bioRxiv (2020).

33. Stoeckius, M. et al. Simultaneous epitope and transcriptome measurement in single cells. Nature methods 14, 865 (2017).

34. Mimitou, E. P. et al. Scalable, multimodal profiling of chromatin accessibility and protein levels in single cells. bioRxiv (2020).

35. Kim, H. J., Lin, Y., Geddes, T. A., Yang, J. Y. H. & Yang, P. CiteFuse enables multi-modal analysis of CITE-seq data. Bioinformatics 36, 4137–4143 (2020).

36. Godfrey, D. I., MacDonald, H. R., Kronenberg, M., Smyth, M. J. & Van Kaer, L. NKT cells: what’s in a name? Nature Reviews Immunology 4, 231–237 (2004).

37. Finak, G. et al. MAST: a flexible statistical framework for assessing transcriptional changes and characterizing heterogeneity in single-cell RNA sequencing data. Genome biology 16, 1–13 (2015).

38. Abdelaal, T. et al. A comparison of automatic cell identification methods for single-cell RNA sequencing data. Genome biology 20, 194 (2019).

39. Maecker, H. T., McCoy, J. P. & Nussenblatt, R. Standardizing immunophenotyping for the human immunology project. Nature Reviews Immunology 12, 191–200 (2012).

