## Supplementary figures and table for "scJoint: transfer learning for data integration of atlas-scale single-cell RNA-seq and ATAC-seq"

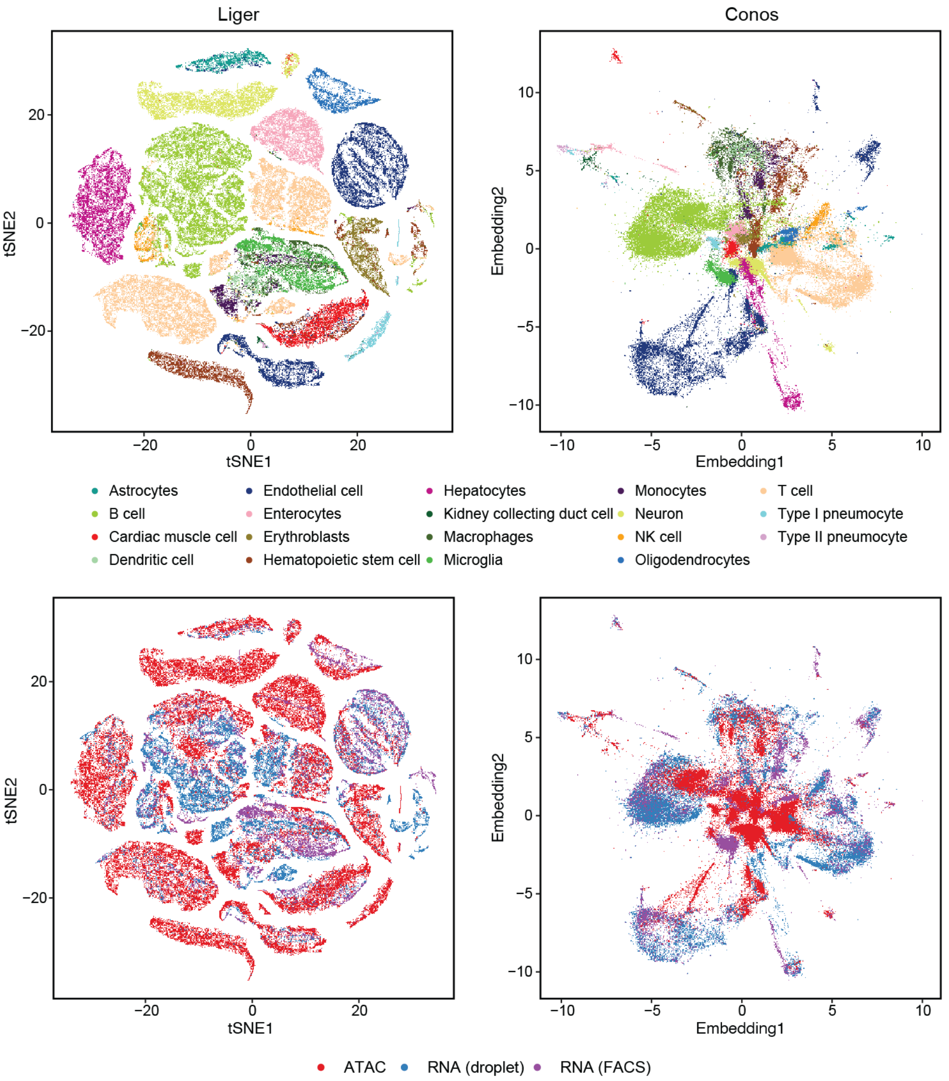


Supplementary Figure S1: tSNE visualization of the overlapping subset data from mouse cell atlases for Liger (first column) and Conos (second column), colored by cell type (first row) and technology (second row).

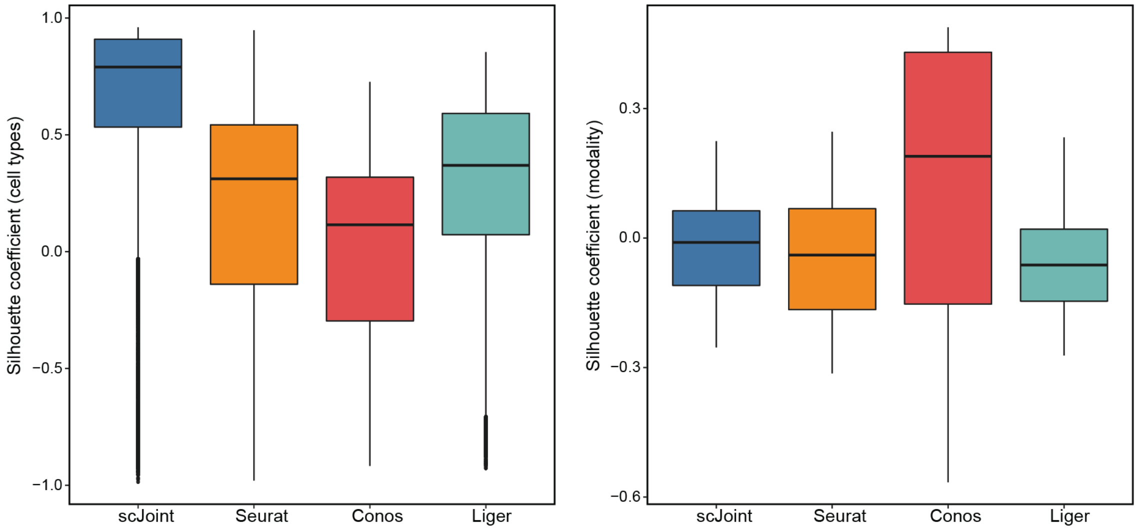


Supplementary Figure S2: Evaluating the joint visualizations of the mouse cell atlas subset data. Boxplots of cell type silhouette coefficients (left) and modality silhouette coefficient (right) for scJoint, Seurat, Conos and Liger.

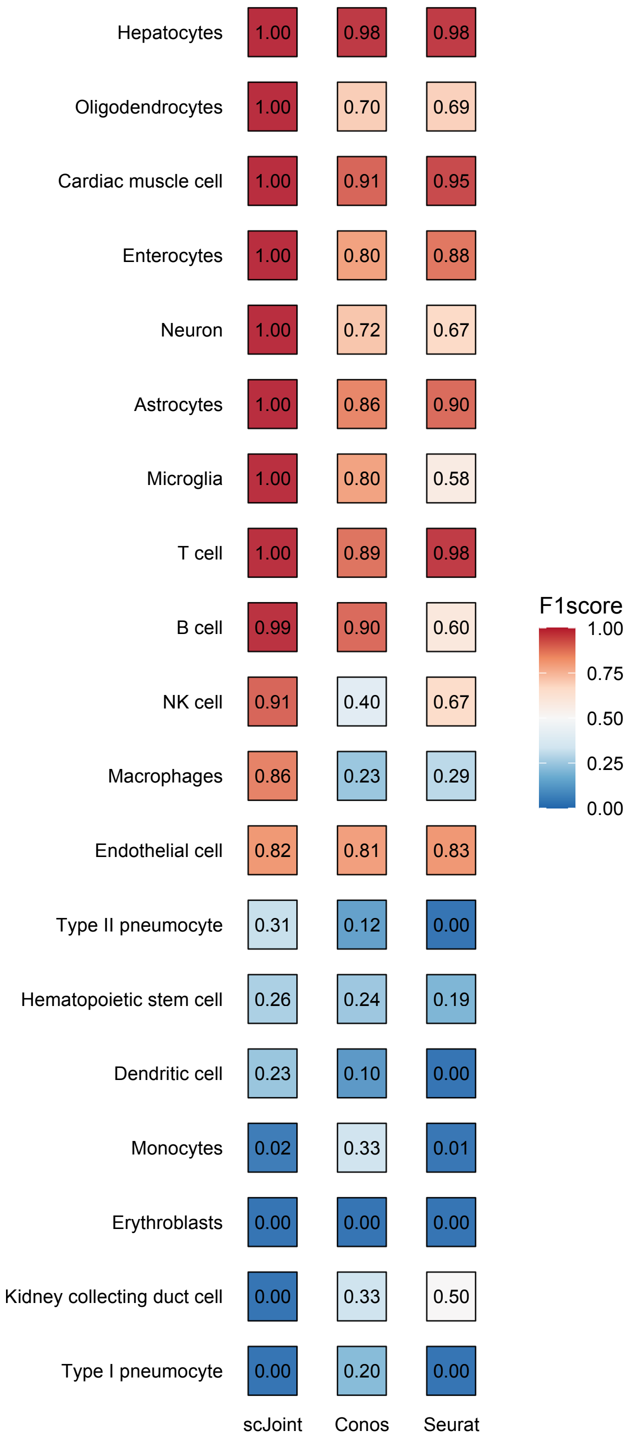


Supplementary Figure S3: Evaluating the accuracy of transferred labels for each cell type in the mouse cell atlas subset data. F1-scores of cell type classification from each method.

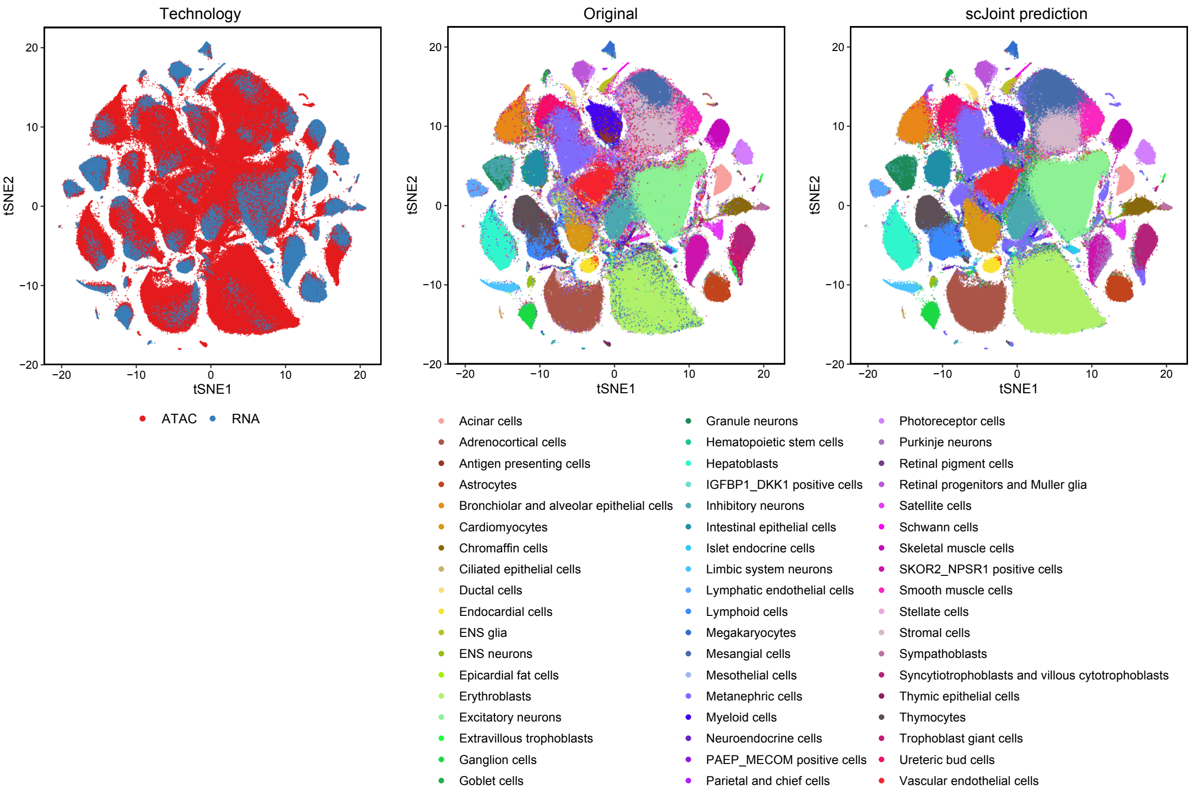


Supplementary Figure S4: tSNE visualization of the overlapping subset data (433,695 cells from scRNA-seq and 656,074 cells from scATAC-seq) from human fetal atlas for scJoint, colored by technology (left), original labels (middle) and scJoint prediction (right).

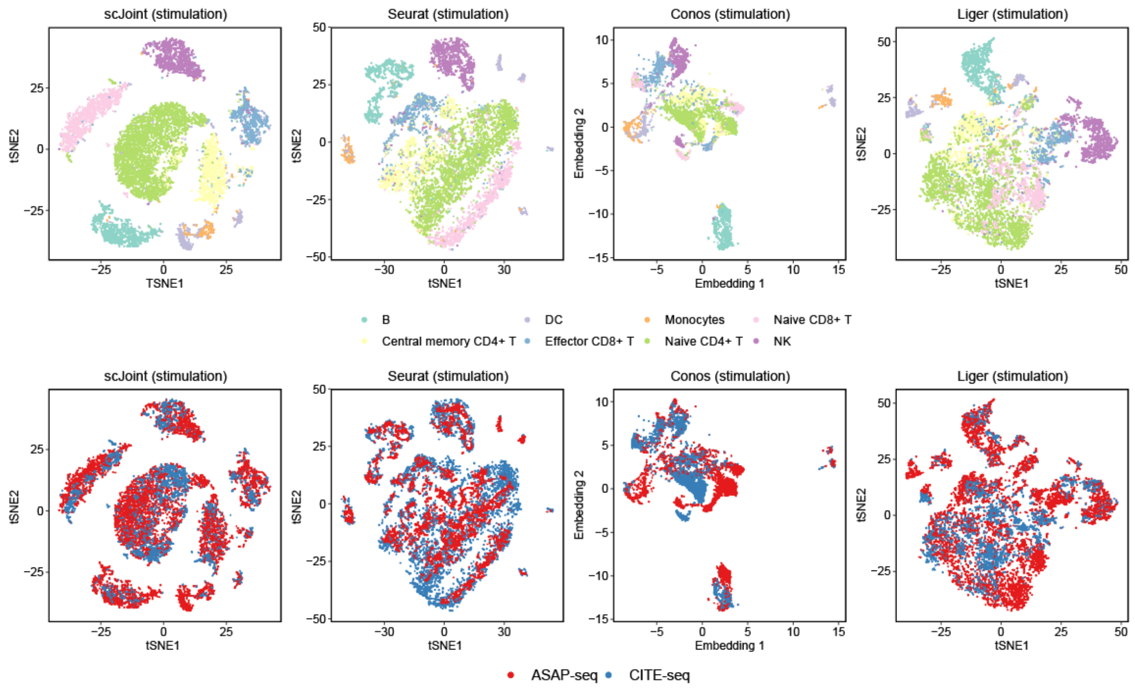


Supplementary Figure S5: tSNE visualization of CITE-seq and ASAP-seq PBMC data under stimulation, generated by scJoint (first column), Seurat (second column), Conos (third column) and Liger (fourth column), colored by cell types (first row) from CiteFuse and manual annotations, and technology (second row).

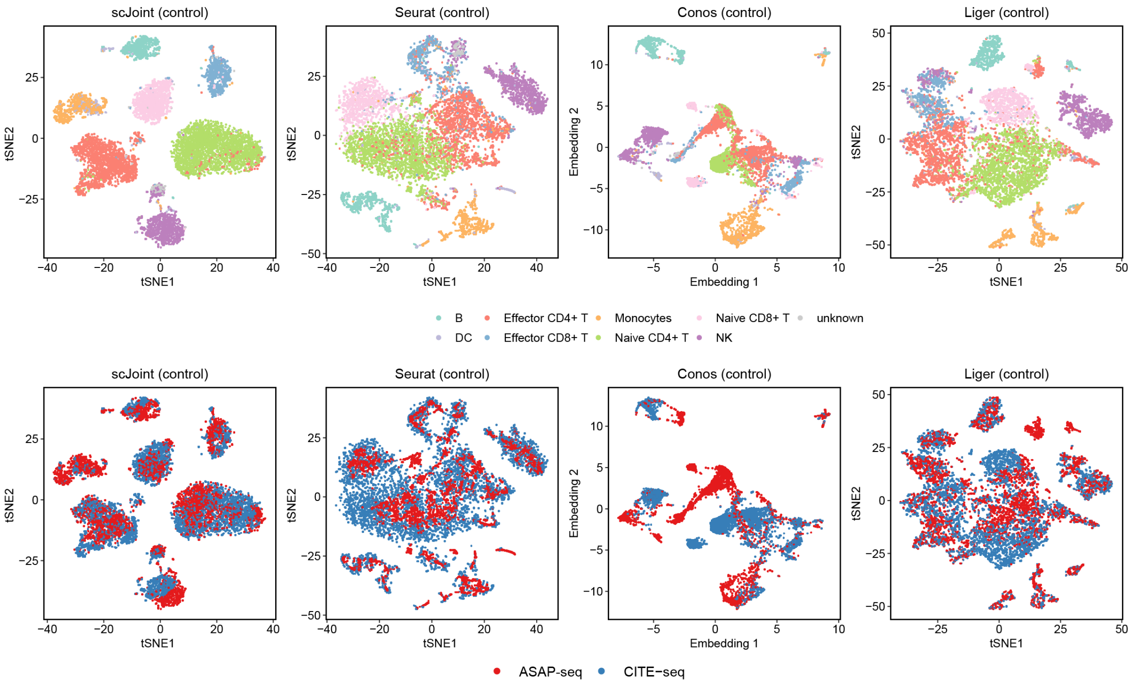


Supplementary Figure S6: tSNE visualization of CITE-seq and ASAP-seq PBMC data under the control condition, generated by scJoint (first column), Seurat (second column), Conos (third column) and Liger (fourth column), colored by cell types (first row) from CiteFuse and manual annotations, and technology (second row).

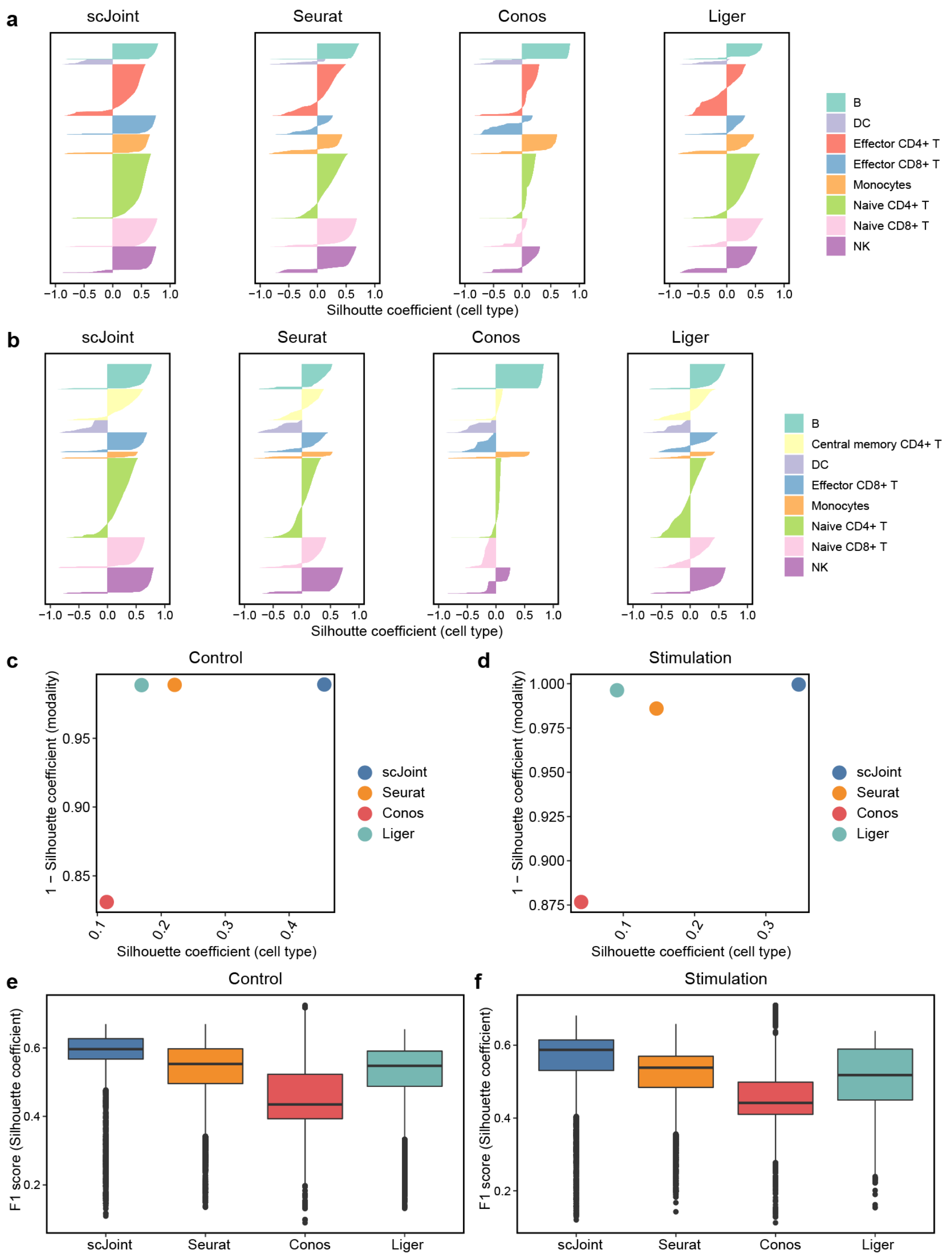


Supplementary Figure S7: Evaluating the joint visualizations of CITE-seq and ASAP-seq PBMC data. (a-b) Barplots of cell type silhouette coefficients for scJoint, Seurat, Conos and Liger for all cells, colored by cell types under (a) control; (b) stimulation. (c-d) Scatter plot of mean silhouette coefficients for scJoint, Seurat, Conos and Liger (left), where the x-axis denotes the mean silhouette coefficients of cell types and the y-axis denotes 1 - mean modality silhouette coefficients under two conditions: (c) control; (d) stimulation; (e-f) Boxplots of F1 scores of silhouette coefficients for scJoint, Liger, Seurat, and Conos, under (e) control and (f) stimulation.

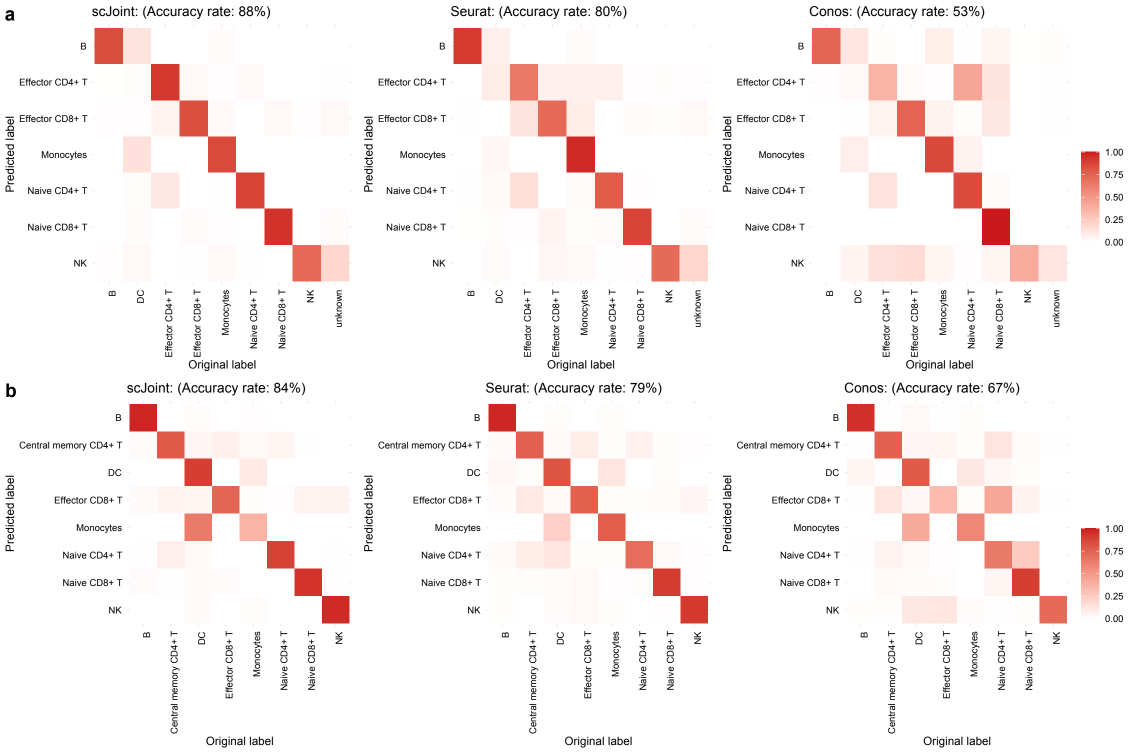


Supplementary Figure S8: Label transfer accuracy in CITE-seq and ASAP-seq PBMC data. Heatmaps show fractions of agreement between the original labels and the transferred labels of scJoint, Seurat and Conos: (a) control; (b) stimulation.

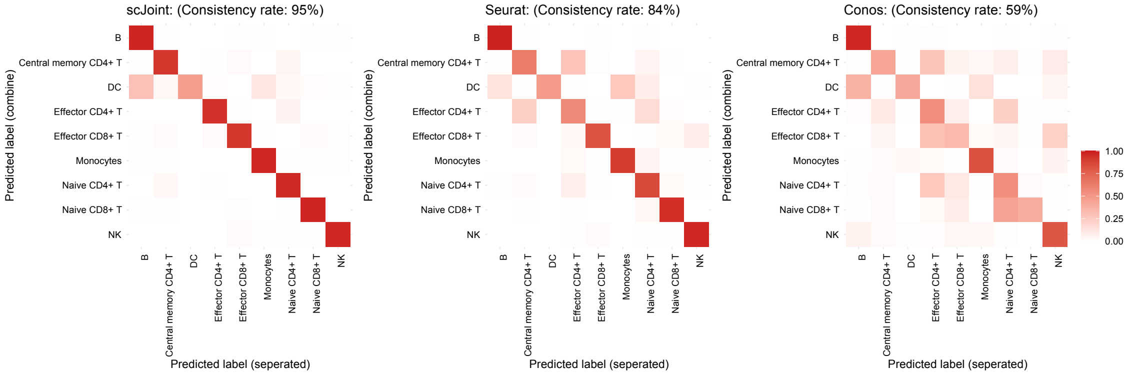


Supplementary Figure S9: Consistency of label transfer in CITE-seq and ASAP-seq PBMC data. Heatmaps show fractions of agreement among the transferred labels from running each method on control / stimulation separately and two conditions jointly: scJoint (left), Seurat (middle) and Conos (right).

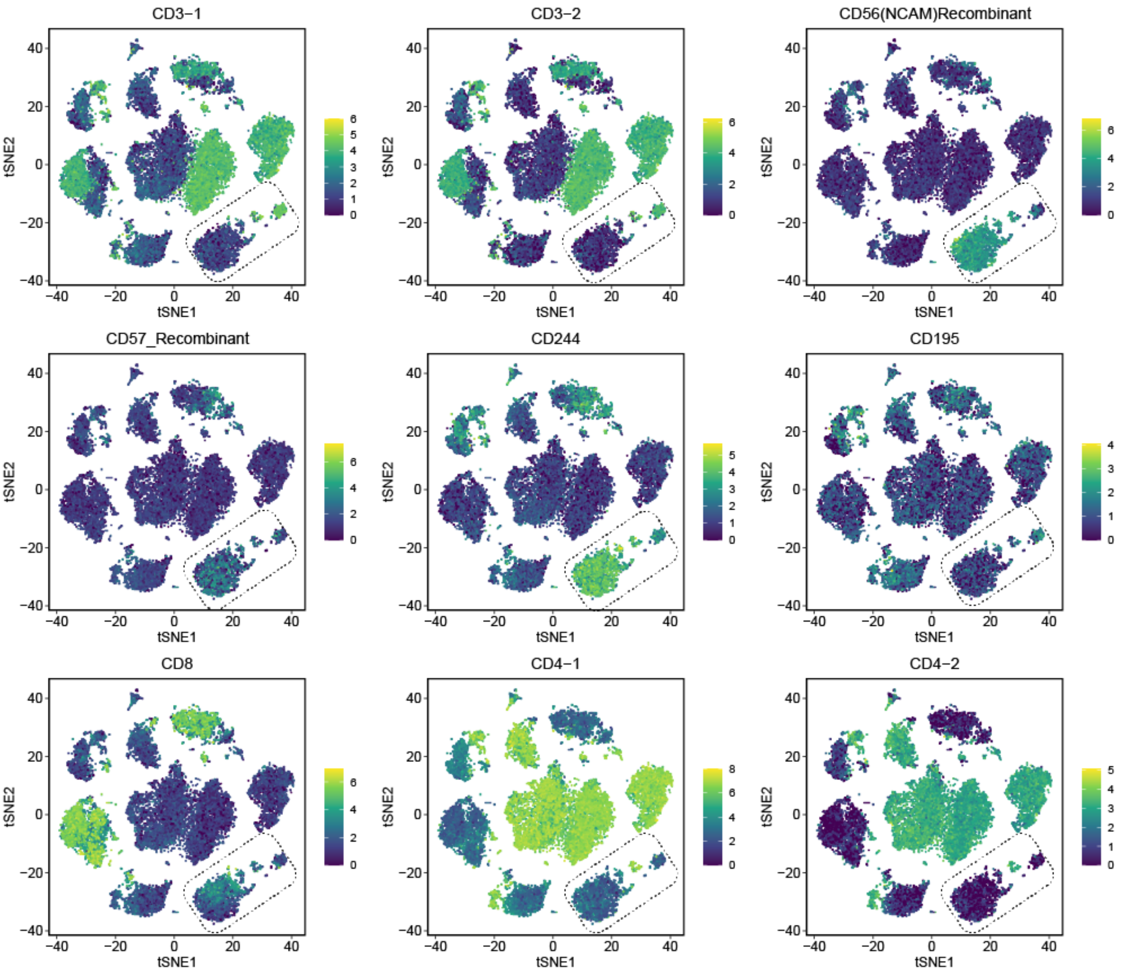


Supplementary Figure S10: ADT expression of NK T cells in CITE-seq and ATAC-seq data.

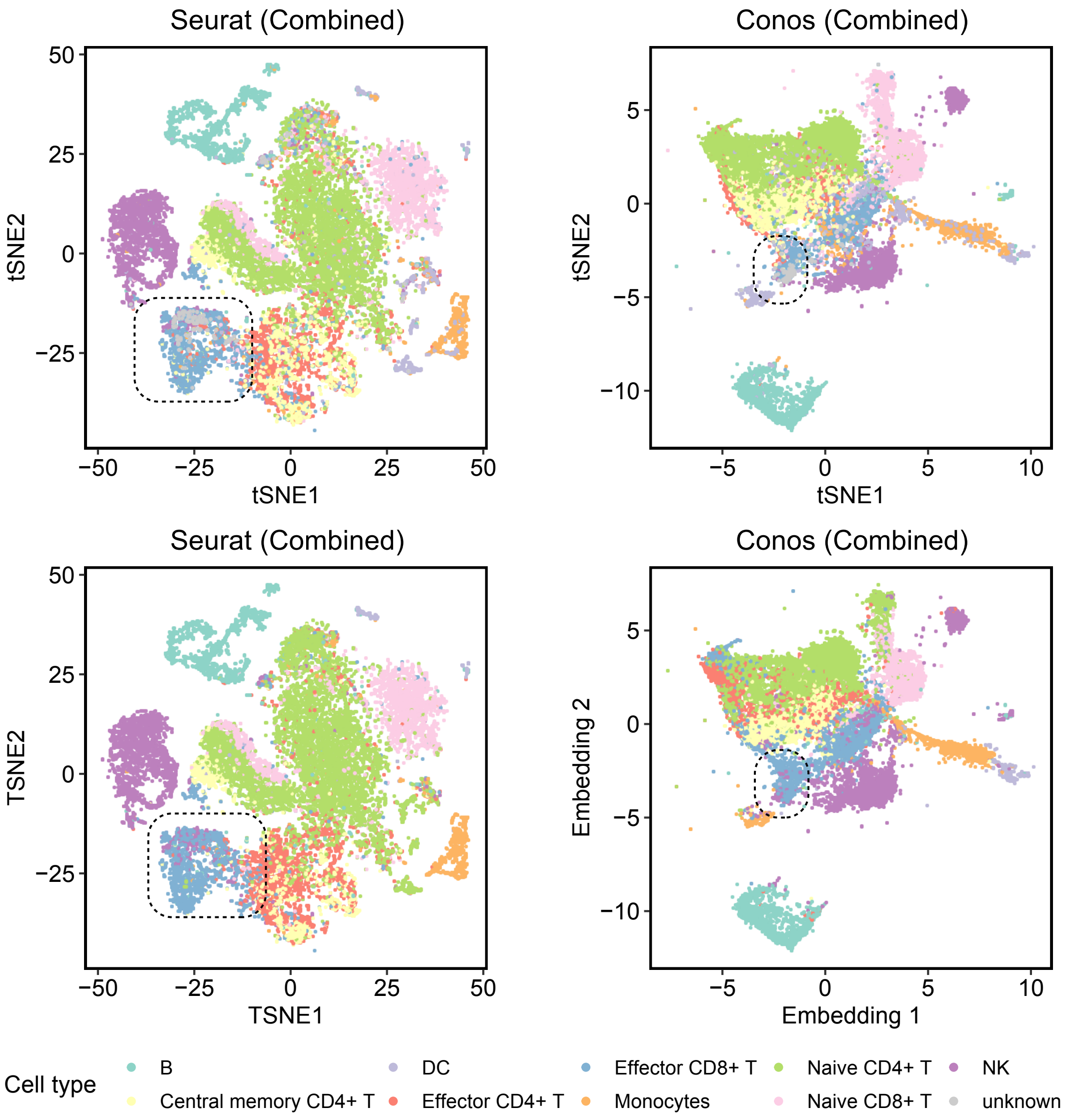


Supplementary Figure S11: tSNE visualization of CITE-seq and ASAP-seq PBMC data under combined conditions, generated by Seurat (first column) and Conos (second column), colored by original cell types (first row) from CiteFuse and manual annotations, and predicted cell types. Cells identified as NK T cells in scJoint visualization are mixed with Effector CD8+ T cells by other methods.

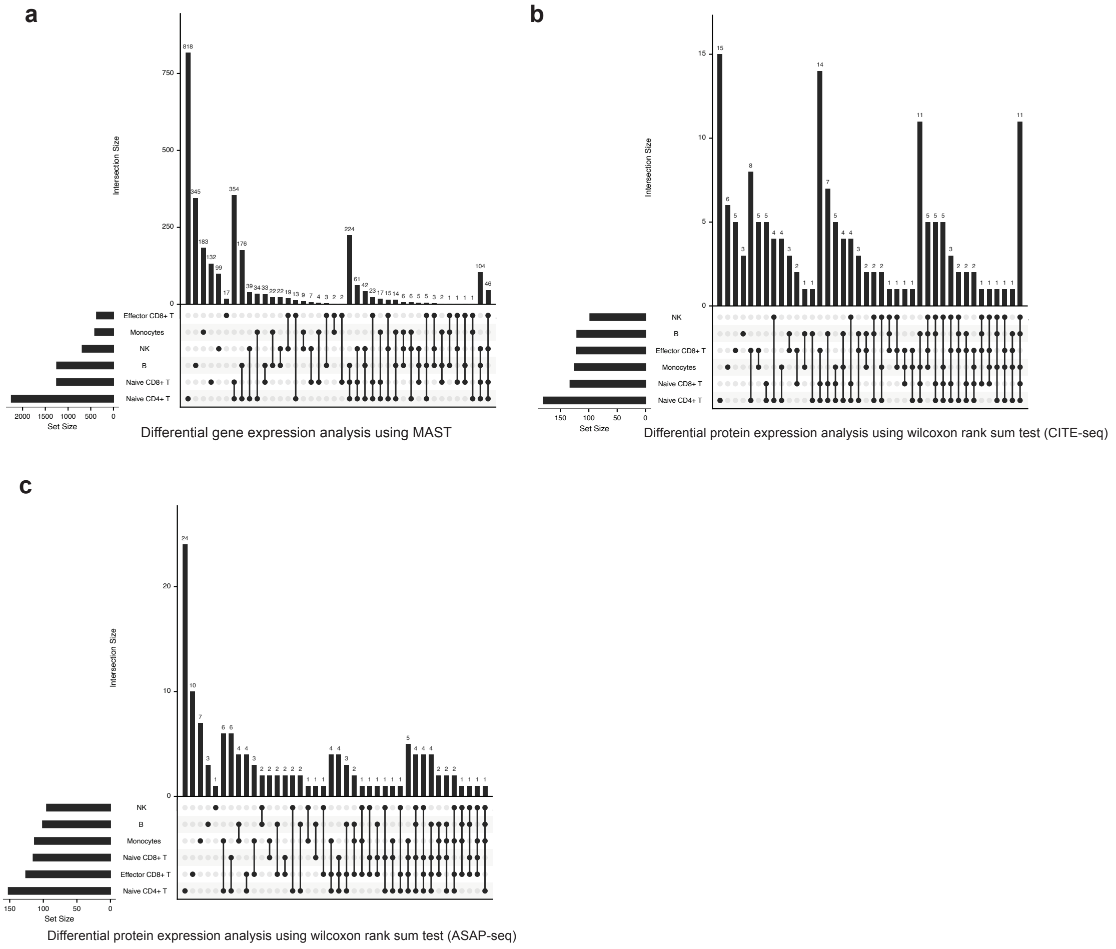


Supplementary Figure S12: Differential expression (DE) analysis across two conditions of CITE-seq and ASAP-seq.

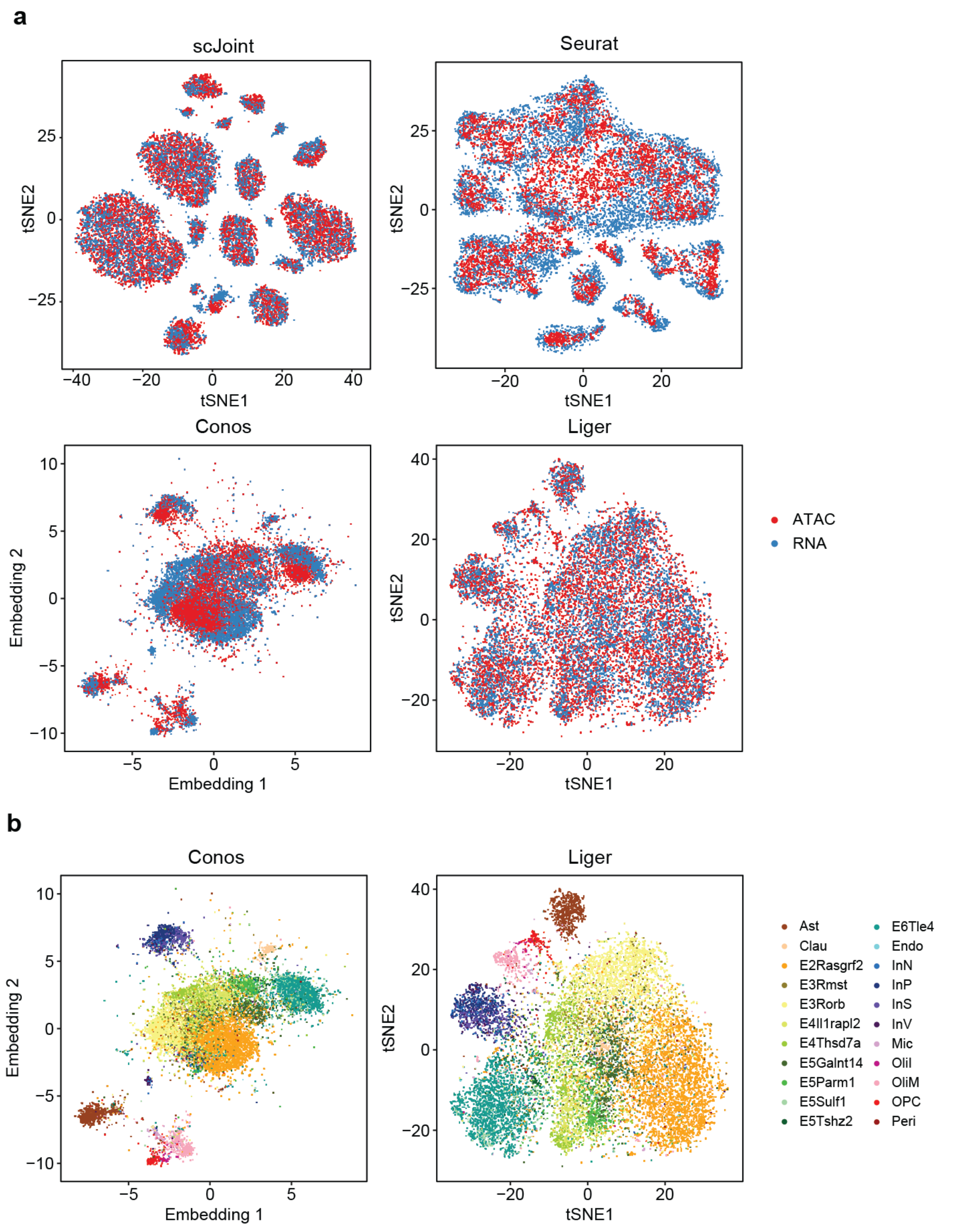


Supplementary Figure S13: (a) tSNE visualization of SNARE-seq data with the RNA and ATAC parts colored separately for unpaired methods: scJoint (top left), Seurat (top right), Conos (bottom left) and Liger (bottom right). (b) tSNE visualization of SNARE-seq data colored by original cell types, generated by Conos (left), Liger (right).

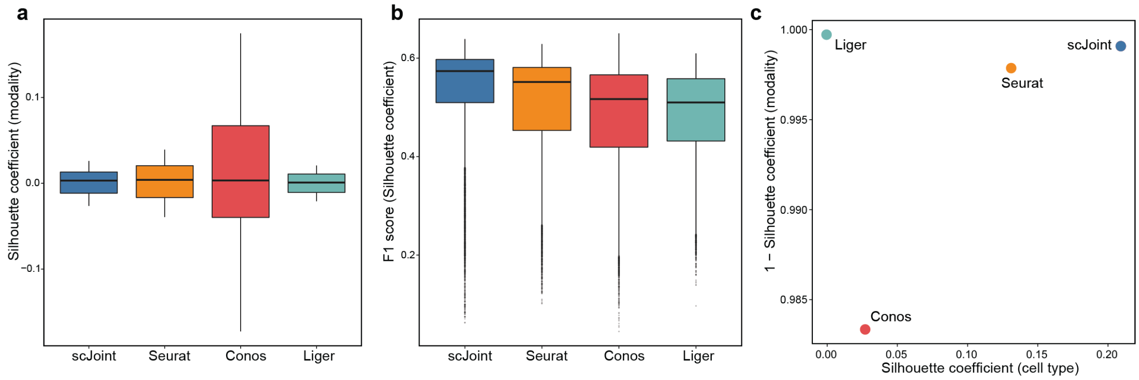


Supplementary Figure S14: Evaluating the joint tSNE visualizations for SNARE-seq data. (a) Boxplots of modality silhouette coefficients for scJoint, Seurat, Conos, and Liger; smaller values indicate better mixing. (b) Boxplots of F1 scores of silhouette coefficients for scJoint, Seurat, Conos, and Liger; larger values indicate better balance. (c) Scatter plot of mean silhouette coefficients for scJoint, Liger, Seurat, and Conos, where the x-axis shows the mean cell type silhouette coefficients and the y-axis shows 1 - mean modality silhouette coefficients; ideal outcomes would lie in the top right corner.

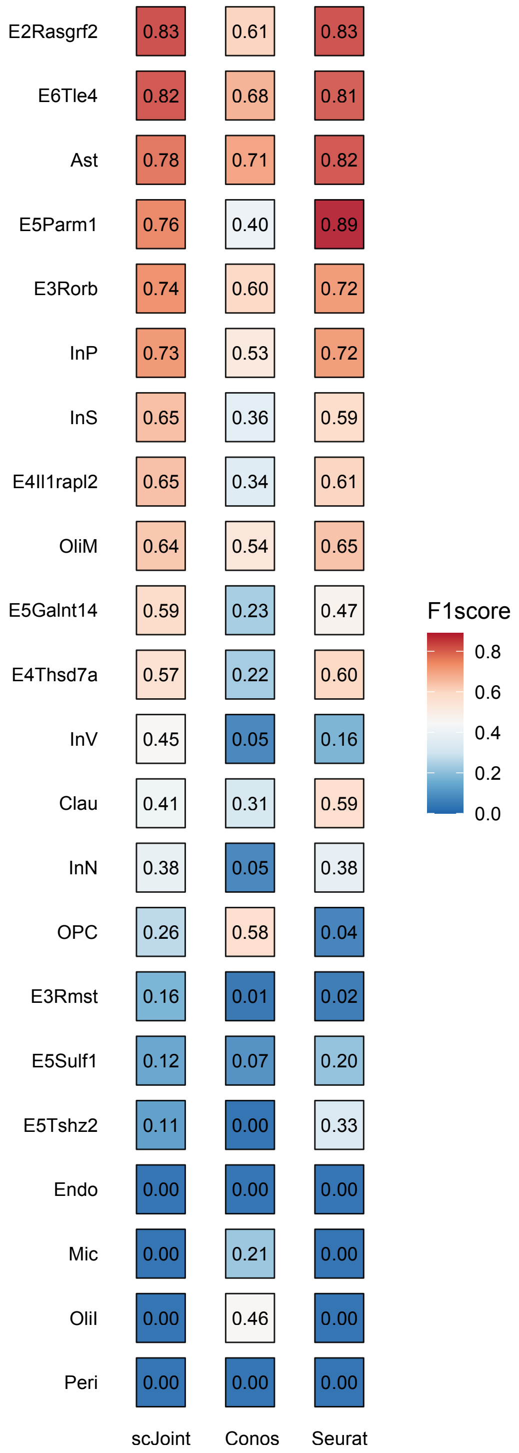


Supplementary Figure S15: Evaluating the accuracy of transferred labels for each cell type in the SNARE-seq data. F1-scores of cell type classification from each method.

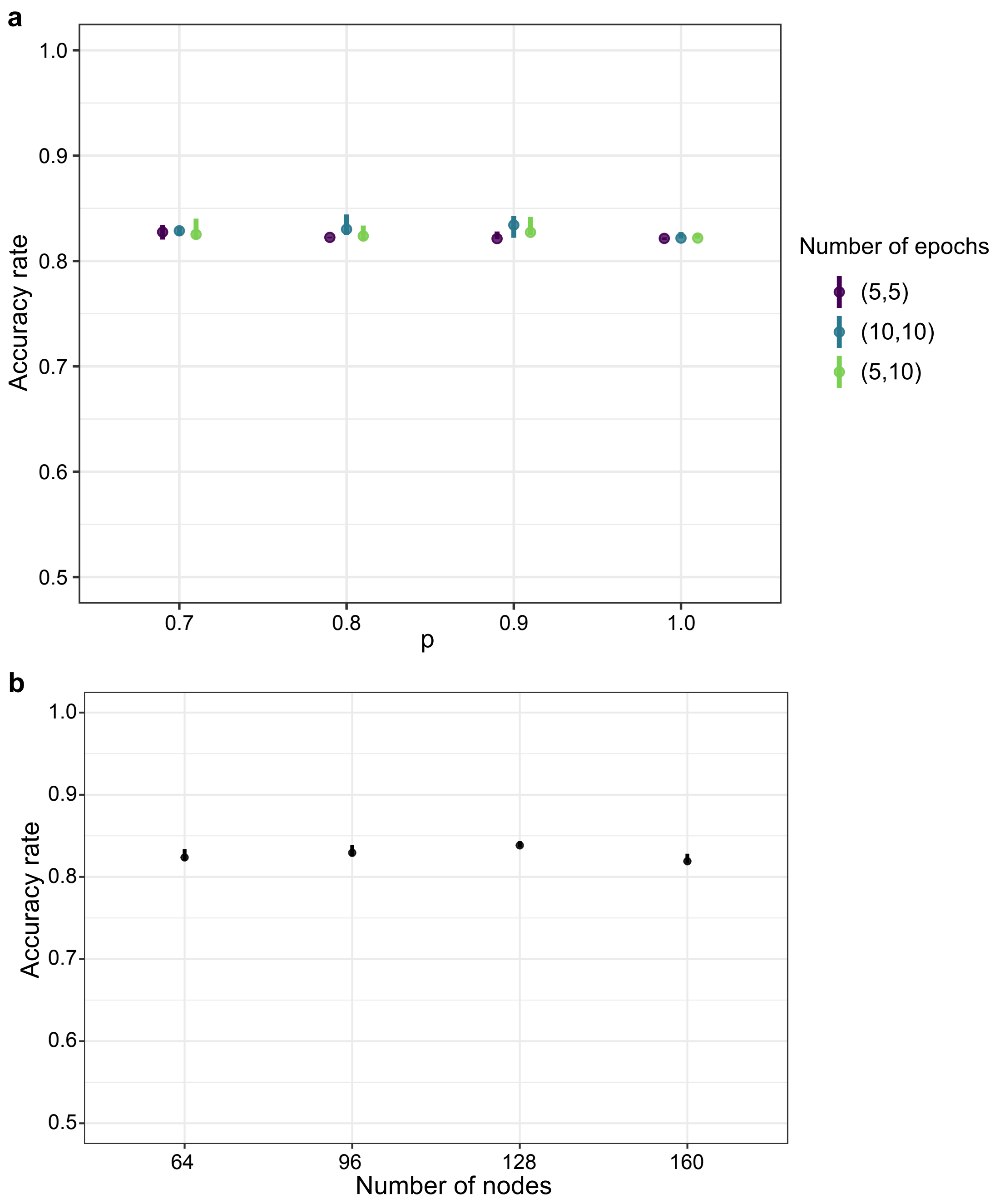


Supplementary Figure S16: Robustness to tuning parameters. Label transfer accuracy of scJoint on the overlapping subset data from the mouse cell atlases when varying (a) the fraction $p$ of data pairs included in the cosine similarity loss ($p = 0.7, 0.8, 0.9, 1.0$) and number of training epochs in Step 1 and 3 ($(5, 5)$, $(10, 10)$, and $(5, 10)$); (b) the number of nodes in the embedding (hidden) layer (number of nodes $= 64, 96, 128, 160$). The dots indicate the medians and the bars indicate the interquartile range from 10 independent runs.

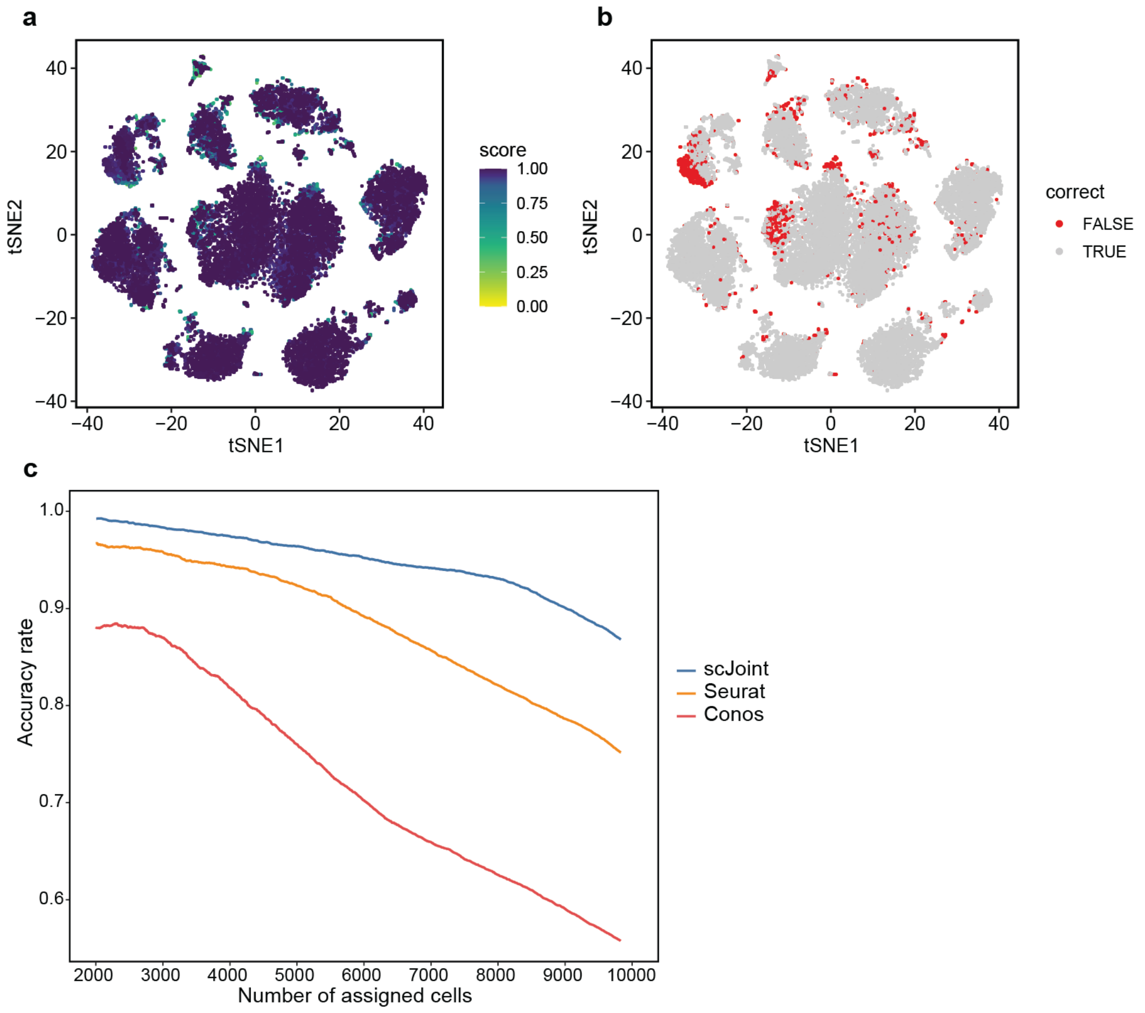


Supplementary Figure S17: Evaluating the probability scores for label transfer in CITE-seq and ASAP-seq PBMC data. (a-b) tSNE visualization using scJoint colored by: (a) probability score of cell type prediction; (b) the correctness of transferred labels. (c) Accuracy rate changes as we change the threshold for probability scores in each method. The x-axis shows the number of cells whose probability scores exceed a given threshold and were assigned a prediction; the y-axis shows the corresponding accuracy rate.

| Dataset | $S$ | $T$ | # epochs (Step 1) | lr (Step 1) | # epochs (Step 3) | lr (Step 3) |
|---|---|---|---|---|---|---|
| Mouse atlas full | 2 | 1 | 5 | 0.01 | 10 | 0.001 |
| Mouse atlas subset | 2 | 1 | 5 | 0.01 | 10 | 0.001 |
| SNARE-seq | 1 | 1 | 10 | 0.01 | 10 | 0.01 |
| Multi-modal control | 1 | 1 | 10 | 0.01 | 10 | 0.01 |
| Multi-modal stimulation | 1 | 1 | 10 | 0.01 | 10 | 0.001 |
| Multi-modal combined | 1 | 1 | 10 | 0.01 | 10 | 0.01 |

Table S1: Training details for each data listing the number of scRNA-seq datasets ($S$), number of scATAC-seq datasets (T), learning rate (lr) and number of training epochs used in Step 1 and Step 3.

| Number of cells | batch size | # epochs (Step 1) | lr (Step 1) | # epochs (Step 3) | lr (Step 3) |
|---|---|---|---|---|---|
| $\leq$ 50k | 256 | 10 | 0.01 | 10 | 0.01 |
| 50k - 500k | 512 | 10 | 0.01 | 10 | 0.01 |
| $\geq$ 500k | 1024 | 10 | 0.01 | 10 | 0.01 |

Table S2: Training details for the human fetal atlas data, including the batch size, learning rate (lr) and number of training epochs used in Step 1 and Step 3.
